# Stress induced TDP-43 mobility loss independent of stress granules

**DOI:** 10.1101/2022.02.28.482242

**Authors:** Lisa Streit, Timo Kuhn, Thomas Vomhof, Albert C. Ludolph, Jochen H. Weishaupt, J. Christof M. Gebhardt, Jens Michaelis, Karin M. Danzer

**Author notes:** <u>Corresponding authors</u>.

## Abstract

TAR DNA binding protein 43 (TDP-43) is closely related to the pathogenesis of amyotrophic lateral sclerosis (ALS) and translocates to stress granules (SGs). The role of SGs as aggregation-promoting “bioreactors” for TDP-43, however, is still under debate. We analyzed TDP-43 mobility and localization under different stress and recovery conditions using live cell single-molecule tracking and super-resolution microscopy. Besides reduced mobility within SGs, a stress induced decrease of TDP-43 mobility in the cytoplasm and the nucleus was observed. Stress removal led to a recovery of TDP-43 mobility, which strongly depended on the stress duration. ‘Stimulated-emission depletion microscopy’ (STED) and ‘tracking and localization microscopy’ (TALM) revealed not only TDP-43 substructures within stress granules but also numerous patches of slow TDP-43 species throughout the cytoplasm. The data provide new insights into the aggregation of TDP-43 in living cells and provide evidence suggesting that TDP-43 oligomerization takes place in the cytoplasm separate from SGs.

## Introduction

TAR DNA binding protein 43 (TDP-43) is a predominantly nuclear DNA-/RNA-binding protein. Cytoplasmic TDP-43 inclusions are a neuropathological hallmark of amyotrophic lateral sclerosis (ALS) and frontotemporal lobar degeneration (FTD) (Neumann et al. 2006; Arai et al. 2006). ALS is a fatal neurodegenerative disease involving the progressive degeneration of the upper and lower motor neurons, leading to paralysis and death within 3 – 5 years after symptom onset (Masrori and Van Damme 2020). FTD is an entity that is clinically and genetically linked to ALS and most often associated with behavioral alterations, personality changes and dementia. Besides the formation of pathological TDP-43 in many FTD and >95% of ALS patients (Neumann et al. 2006; Arai et al. 2006), also the existence of ALS-causing TDP-43 mutations attributes and important role of TDP-43 for the pathogenesis of ALS/FTD (Kabashi et al. 2008).

Understanding the aggregation process is therefore broadly investigated and highly relevant for understanding the molecular pathogenesis of ALS/FTD. One starting point for the aggregation of TDP-43 could be its localization to and interaction with stress granules (SGs). SGs constitute a small reaction volume, densely packed with aggregation prone proteins and have been suspected to serve as a starting point for aggregation of several proteins related to neurodegeneration (Baradaran-Heravi, Van Broeckhoven, and van der Zee 2020; Wolozin and Ivanov 2019; Jeon and Lee 2021). However, it is still under debate, whether TDP-43 localization to stress granules prevents or stimulates its pathological aggregation. Recent studies hint towards a mechanism where localization to stress granules and RNA interactions may exert, an initially protective rather than a disease promoting role (Fernandes et al. 2020; McGurk et al. 2018; Mann et al. 2019; Gasset-Rosa et al. 2019).

How TDP-43 mobility changes on a molecular level and in different cellular compartments and conditions is not known so far. It will be of central importance to understand if TDP-43 exhibits different mobility regimes depending on the spatial subcellular distribution.

We therefore employed single-molecule live cell tracking, super-resolution stimulated emission depletion (STED) and tracking and localization microscopy (TALM), to gain more insight into TDP-43 behavior and spatial distribution on a molecular level in living cells.

## Results

### Generation of double-transgenic Halo-TDP-43 and G3BP1-SNAP cell lines enables specific and bright labeling for single-molecule tracking experiments

To study TDP-43 aggregation, its mobility in different cellular compartments and its role in stress granules on a single-molecule level, we engineered dual-transgenic TDP-43-Halo and G3BP1-SNAP cell lines (see Methods). As the attachment of bright, membrane permeable and photo-stable fluorophores is a key requisite for single-molecule studies, the commonly used HaloTag (Los et al. 2008) and SNAP-Tag (Keppler et al. 2004) systems were used in this study. To ensure that the fusion of the HaloTag (33 kDa) to TDP-43 does not alter the protein’s localization and function, the HaloTag was fused to either the N- or the C-terminus of TDP-43, named ^Halo^TDP-43 and TDP-43^Halo^, respectively (Fig. 1A, S1A). To spatially assign stress granules during image analysis we used G3BP1^SNAP^ as a stress granule marker.

**Figure 1:**
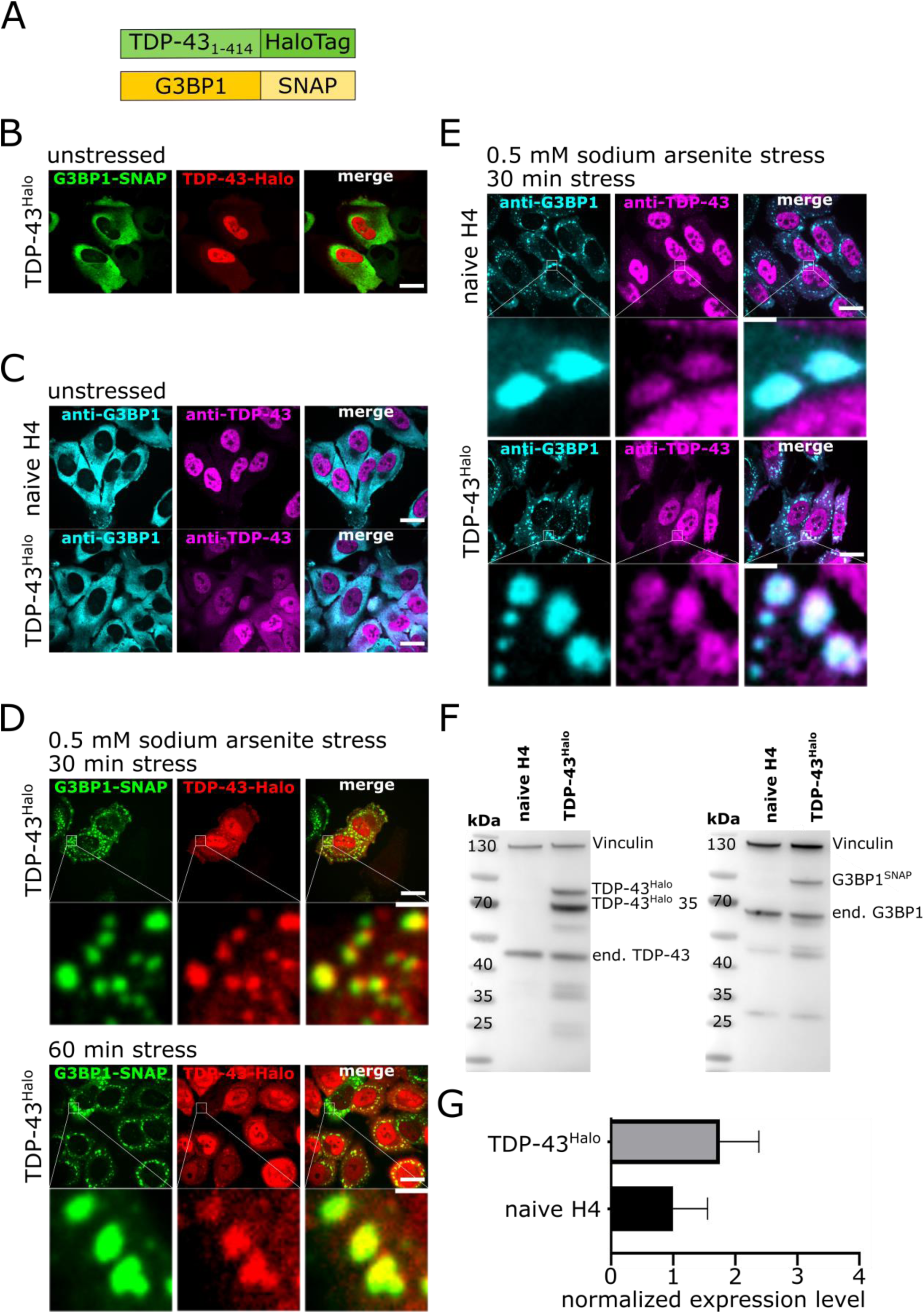
Generation and quality control of TDP-43^Halo^ cell line. A. Schematic overview of the TDP-43^Halo^ and G3BP1^SNAP^ constructs. B. Spinning disk confocal images of the TDP-43^Halo^ cell line under unstressed conditions (red: TDP-43-TMR, green: G3BP-SiR, scale bar 20 µm). C. Spinning disk confocal images of immune-labelled naïve H4 cells as well as TDP-43^Halo^ cells under unstressed conditions (cyan: anti-G3BP1-Alexa532, magenta: anti-TDP-43-Alexa647, scale bar 20 µm). D. Spinning disk confocal images of the TDP-43^Halo^ cell line under 30 min and 60 min sodium arsenite treatment (red: TDP-43-TMR, green: G3BP-SiR, scale bar 20 µm and 2 µm). E. Spinning disk confocal images of the immunostained TDP-43^Halo^ cell line and naïve H4 cells under 60 min sodium arsenite treatment (magenta: anti-TDP-43-Alexa647, cyan: anti-G3BP-Alexa532, scale bar 20 µm and 2 µm). F. Western Blot overview of the TDP-43^Halo^ cell line and naïve H4 cells stained with anti-vinculin, anti-TDP-43 or anti-G3BP1 antibodies showing proper expression of the transgenic constructs. G. Quantification of the overexpression for the TDP-43^Halo^ cell line compared to endogenous TDP-43 in naïve H4 cells shows a 1.5 – 2x overexpression of TDP-43^Halo^.

We first ensured proper functionality of the tagged TDP-43 and G3BP1 constructs in the transgenic cell lines by Sir-Halo and TMR-SNAP staining, as well as immunostaining of endogenous proteins, and compared observed localizations as a function of stress duration to that in naïve H4 cells (Fig. 1B, C). In contrast to a diffusive cytoplasmic G3BP1 signal under unstressed conditions, G3BP1 positive stress granules containing TDP43^Halo^ could be observed after 30 min or 60 min of sodium arsenite stress (Fig. 1D), which was also confirmed by immunostaining (Fig. 1E).

We then quantified the total of TDP-43^Halo^ and G3BP1^SNAP^ overexpressed proteins by Western blot analysis (Fig. 1F). Densitometric analysis showed an over-expression of 1.5 – 2 fold for the TDP-43^Halo^ construct compared to endogenous TDP-43 in naïve H4 cells (Fig. 1G). We focused on C-terminally tagged TDP-43^Halo^, since it allows for the visualization of both, full-length and fragmented TDP-43 species (Fig. 1F). Complementary data for the N-terminally tagged TDP-43 construct (^Halo^TDP-43) are given in the supplementary materials (Fig. S1, S2).

C-terminally tagged TDP-43 shows formation of a prominent 35 kDa TDP-43 fragment and an increased cytoplasmic localization, which is not seen for the N-terminally tagged TDP-43. Fragmentation of TDP-43 leads to the disruption or complete abolishment of the nuclear localization sequence (NLS) (François-Moutal et al. 2019; Berning and Walker 2019). A disrupted or missing NLS leads to a subsequent accumulation of TDP-43 fragments in the cytoplasm, and could explain the observed slightly higher cytoplasmic level of TDP-43^Halo^ as compared to the naïve H4 cells (Fig. 1C).

Taken together, the attachment of either Tag the respective protein, does not alter the formation of stress-induced phase-separated compartments or their interaction of G3BP1 and TDP-43 with latter. This establishes the transgenic cell lines as a new model system for studying stress induced dynamical changes of TDP-43 mobility using single-molecule imaging of Halo-tagged TDP-43 within different cellular compartments.

### Single-molecule tracking monitors TDP-43 movements in the nucleus, the cytoplasm and stress granules in living cells

To monitor TDP-43 in a region specific manner, we established a single molecule tracking pipeline, using photoactivatable Janelia Fluor 646 for labelling of Halo-tagged TDP-43 (Grimm et al. 2016) and TMR-labelling for G3BP1^SNAP^ (Figure 2A). The usage of a photoactivatable dye enables labelling at high molar concentrations (mM range) and single-molecule tracking concentrations are achieved by continuously activating only a small subset of labelled TDP-43^Halo^ per frame. Thus, a high number of frames can be recorded while continuously controlling the emitter density per frame (Hansen et al. 2020; Niewidok et al. 2018).

**Figure 2:**
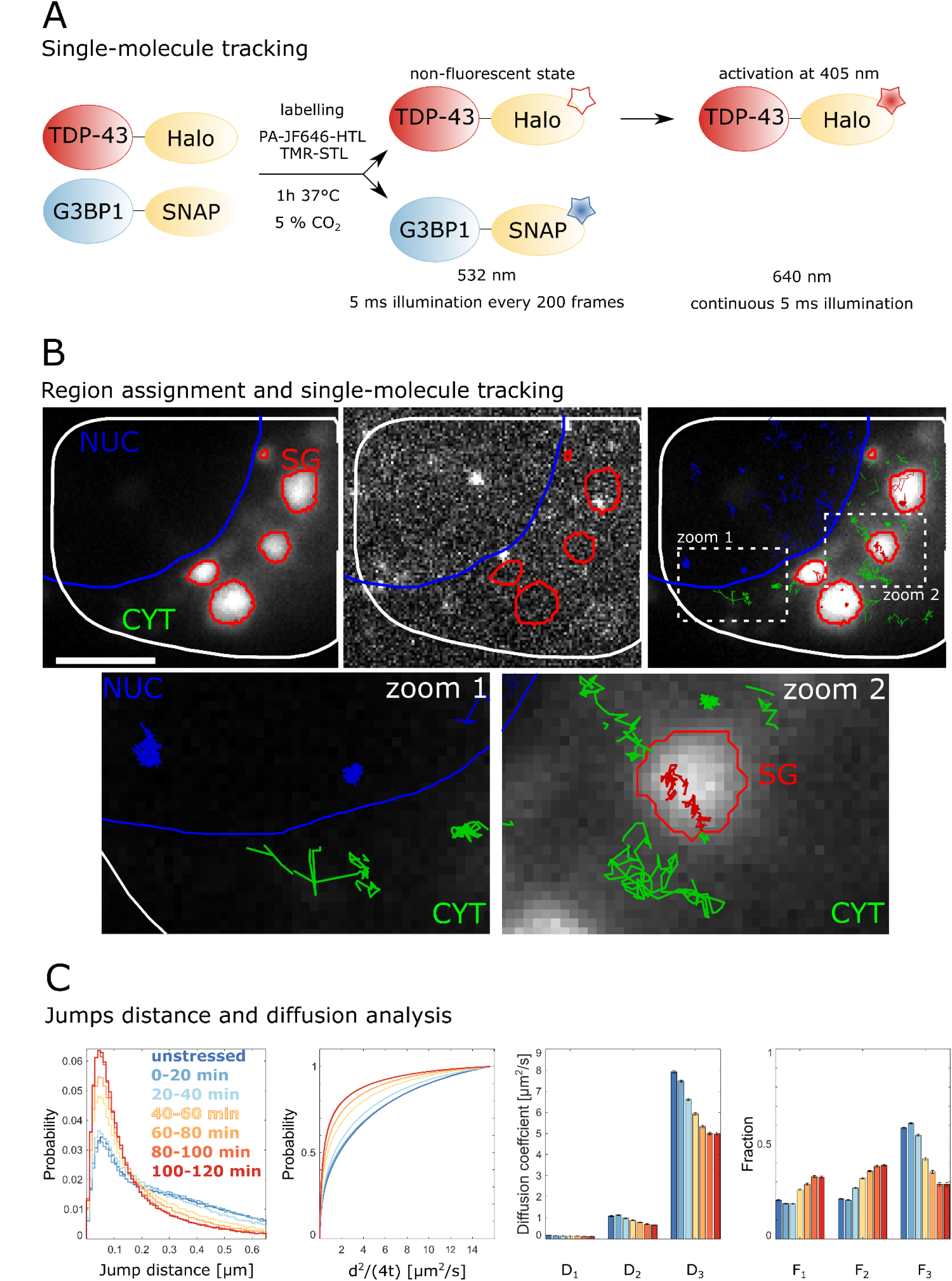
Overview of the single molecule tracking experiments and diffusion analysis. A. Labelling and illumination scheme. TDP-43^Halo^ and G3BP1^SNAP^ are labelled with PA-JF646-HaloTag ligand (HTL) and TMR-SNAP-Tag ligand (STL) for 1h at 37°C and 5% CO_2_ 24 hours before imaging. TDP-43^Halo^ is illuminated continuously with 640 nm and the PA-JF646-dye is continuously activated with 405 nm, the TMR-dye is illuminated with 532 nm and imaged every 200 frames. B. Overview of the region assignment and region-specific track assignment (cytoplasm/tracking region: white outline and green tracks; nucleus: blue outline and tracks; stress granules: red outline and tracks). Zoom-ins and exemplary tracks for all regions are shown and show both, mobile TDP-43^Halo^ as well as more immobile molecules. Scale bar 5 µm. C. Jump distance analysis and 3-exponential fitting results giving three different diffusion coefficients and fractions slow (D_1_/F_1_), medium (D_2_/F_2_) and fast (D_3_/F_3_), in a stress-time-course experiment of TDP-43^Halo^ cells. Stress duration increased from unstressed (blue) to 120 min (red) of 0.5 mM sodium arsenite stress.

TDP-43^Halo-PA-JF646^ molecules were imaged for 120 min under unstressed or stressed conditions (0.5 mM sodium arsenite) using continuous 405 nm activation. In addition, every 200 frames (i.e. every 1.34 s) the G3BP1^SNAP-TMR^ channel was recorded. The movies were grouped into 20 min time slots and tracking and diffusion analysis was performed (Material and Methods). Figure 2B shows the general experimental workflow for region assignment and tracking analysis. Fluorescence from the G3BP1^SNAP^ constructs was used in a control channel to assign cellular regions for analysis. The cellular outline (white line) and the nuclear outline (blue line) are drawn manually (see Materials and Methods). The stress granules (red line) are assigned by an intensity threshold of the G3BP1^SNAP^ signal and tracked dynamically over the whole movie. In each frame, single TDP-43^Halo^ molecules were localized and their position was linked through successive frames of the movie, yielding single-molecule tracks (Figure 2B). Tracks crossing from one region to another (e.g. from the cytoplasm to stress granules) are split at the region border and the parts of the tracks are assigned to the respective region.

Next, displacement and cumulative displacement histograms are computed from all jumps within the tracks subjected to analysis. To reduce the bias towards slower moving and bound molecules, only the first 5 jumps of every tracked single-molecule time-trace were used for further analysis (Hansen et al. 2018). Diffusion coefficients and the respective fractions can be computed by fitting the cumulative displacement histograms with a multi-exponential fit function (Kuhn et al. 2021; Mazza et al. 2012, see Material and Methods). For the extraction of the apparent diffusion coefficients, a three-exponential fit function was used, since it fitted best (compared to mono- and double exponential) to the cumulative jump-distance histogram (Supplementary fig. S3). This results in three diffusion regimes (D_1_/slow, D_2_/medium, D_3_/fast). The three fractions (F_1_/slow, F_2_/medium, F_3_/fast) indicate the proportion of TDP-43^Halo^ molecules in each diffusion regime. An overview of the fitting parameters and fit quality is given in supplementary figure S3. Three diffusion coefficients allow to account not only for the assessment of a bound and mobile fraction but also for the aspect of anomalous diffusion (Guigas, Kalla, and Weiss 2007). Figure 2C depicts the cumulative jump distance histograms computed from detected TDP-43^Halo^ tracks, recorded under unstressed and different stress conditions. These data will be discussed in more detail in the following section (Figure 3). For simplicity the effective diffusion coefficient D_eff_ can be computed (see Material and Methods), representing the weighted average of all diffusion coefficients and respective fractions.

**Figure 3:**
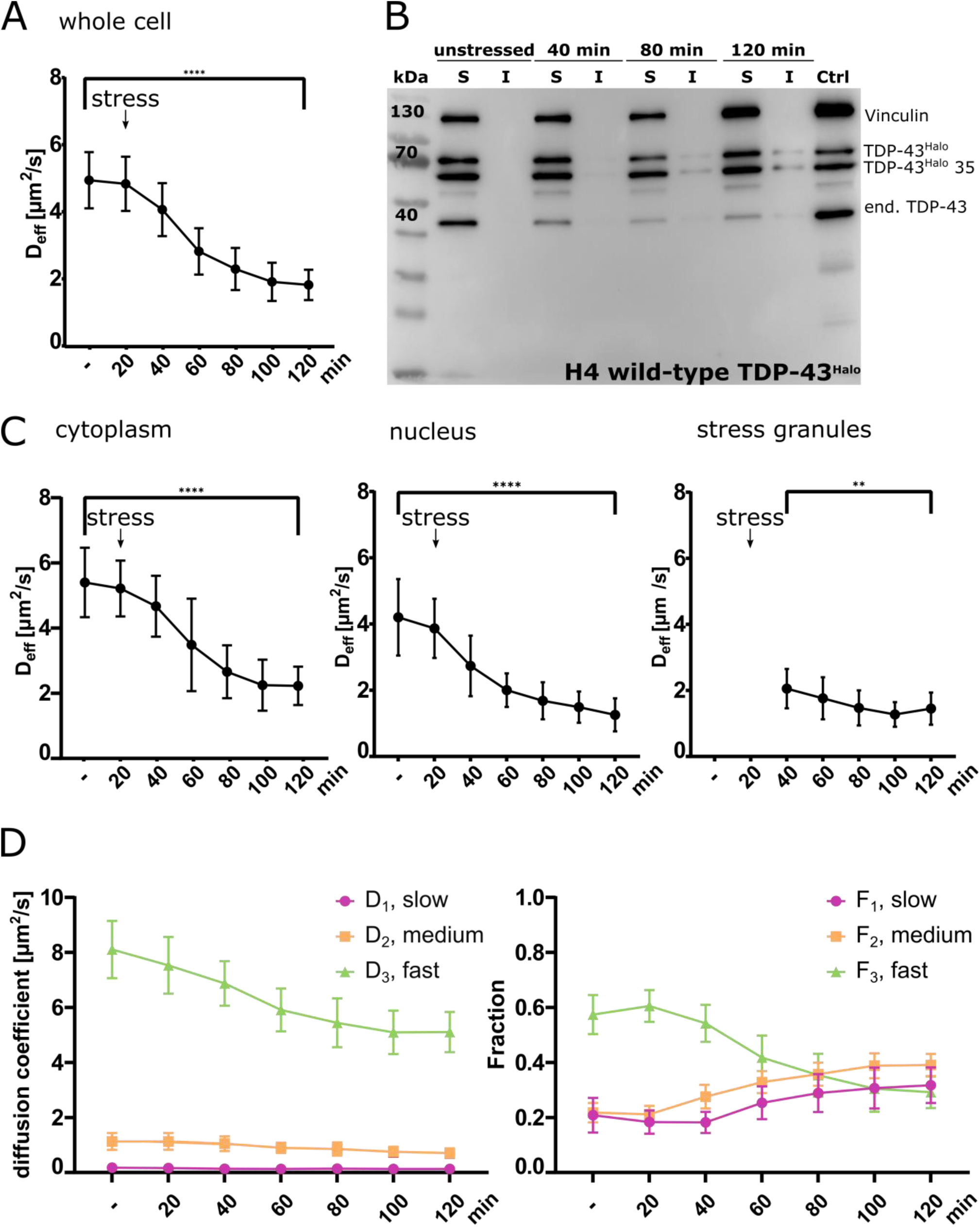
Sodium arsenite stress leads to a reduction of TDP-43^Halo^ mobility in specific cellular regions (cytoplasm, nucleus, stress granules). A. Stress time course of the effective diffusion coefficient D_eff_ for the TDP-43^Halo^ construct (whole cell, mean + STD, Welch’s t-test). B. Solubility Assay of the TDP-43 construct. Solubility was assessed under unstressed conditions and different stress time-points. An increasing insoluble TDP-43 fraction was observed with increasing stress duration (unstressed, 40 min, 80 min and 120 min of 0.5 mM sodium arsenite treatment, anti-Vinculin, anti-TDP-43). C. Stress time course of the effective diffusion coefficient D_eff_ for the TDP-43^Halo^ construct (cytoplasm, nucleus, stress granule, mean + STD, Welch’s t-test). D. Analysis of the different diffusion constants (slow/D_1_/magenta, medium/D_2_/orange, fast/D_3_/green) and the respective fraction within the stress time-course experiment (whole cell). For all experiments, standard deviations were calculated from the movie-wise distribution of the plotted value and statistical significance was assessed with a multiple unpaired t-test with Welch’s correction. P-value ranges: <0.0001 ****, 0.0002 ***, 0.0021 **, 0.032 *, 0.123 ns

In summary, the imaging and analysis pipeline established here, provides a platform to assess TDP-43 mobility in different subcellular regions and to investigate TDP-43 diffusion behavior under stressed and unstressed conditions.

### Sodium arsenite exposure reduces TDP-43 mobility not only in stress granules but also in the nucleus and cytoplasm

To investigate the region specific mobility changes of TDP-43 under sodium arsenite stress, single-molecule tracking analysis was employed (Fig. 3). To biochemically validate the nature of TDP-43 slow-down, additionally, the solubility of *TDP-43*^*Halo*^ and endogenous TDP-43 was assessed using a solubility assay and subsequent Western Blotting.

At first, the effective diffusion coefficient D_eff_ of TDP-43^Halo^ was determined for the whole cell under different stress conditions (Fig. 3). After 120 min of sodium arsenite stress, *TDP-43*^*Halo*^ shows an overall significant reduction in mobility as compared to unstressed conditions (unstressed: D_eff_ = 4.94 µm^2^/s, standard deviation: 0.84 µm^2^/s, stressed: D_eff_ = 1.82 µm^2^/s, standard deviation: 0.45 µm^2^/s, statistical test: Welch’s t-test), suggesting oligomerization of *TDP-43*^*Halo*^ or localization to small compartments that restrict mobility (Fig. 3A). As shown in Figure 3B, an increase of the insoluble TDP-43^Halo^ fraction was observed with increasing stress, starting between 40 and 80 min after stress onset, although the band is rather faint as compared to the soluble fraction. Under unstressed conditions, however, all TDP-43 species are found in the soluble fraction.

When comparing these data to the single molecule tracking data it is likely that the reduced mobility of TDP-43^Halo^ with increasing stress is partly caused by the formation of insoluble TDP-43 aggregates. The reduction of D_eff_ is very pronounced, however, the bands depicting the insoluble TDP-43 fraction is rather faint. This indicates an aggregate-independent mechanism of TDP-43 mobility reduction, since the small amount of insoluble TDP-43 detected by the Western blot alone, cannot explain the observed strong reduction in TDP-43 mobility.

To study *TDP-43*^*Halo*^ mobility in different subcellular compartments and to identify the contribution of different subcellular fractions of TDP-43^Halo^ to the overall mobility reduction seen in the whole cell, diffusion data were analyzed in a region-specific manner.

TDP-43^Halo^ shows the highest mobility in the cytoplasm under unstressed conditions with an effective diffusion coefficient of 5.40 µm^2^/s (standard deviation: 1.07 µm^2^/s) and sodium arsenite stress leads to a significant and continuous reduction of the effective diffusion coefficient to 2.23 µm^2^/s (standard deviation: 0.59 µm^2^/s) after 120 min. Since the HaloTag is attached to the C-terminus of TDP-43 and TDP-43 can be fragmented N-terminally (Arai et al. 2010), the construct visualizes both, full-length as well as fragmented TDP-43^Halo^. A high mobility in the cytoplasm might therefore reflect the presence of full-length and fragmented TDP-43. Under unstressed conditions, TDP-43^Halo^ mobility in the nucleus is in general slower as compared to the cytoplasm (D_eff_ = 4.20 µm^2^/s, standard deviation: 1.15 µm^2^/s), indicating an often bound or confined state of TDP-43 in the nucleus (Gasset-Rosa et al. 2019; Ayala et al. 2008). Notably, also in the nucleus D_eff_ reduced continuously with stress duration reaching the lowest value after 120 min with D_eff_ = 1.26 µm^2^/s (standard deviation: 0.50 µm^2^/s, Fig. 3C).

Stress granules are often discussed as the site of TDP-43 aggregation within the cell (Dewey et al. 2012; Chen et al. 2019; Fernandes et al. 2020). To test this hypothesis, we measured stress-dependent mobility of TDP-43^Halo^ within stress granules. First stress granules could be assigned after 20 min of sodium arsenite stress. Although we observed a substantially lower TDP-43^Halo^ mobility in the stress granules first assigned at 20 min (D_eff_ = 2.05 µm^2^/s (standard deviation: 0.60 µm^2^/s), compared to the unstressed value in the cytoplasm (D_eff_ = 4.20 µm^2^/s, standard deviation: 1.15 µm^2^/s), prolonged stress only led to a modest further reduction of mobility within stress granules reaching at 120 min a value of D_eff_ = 1.45 µm^2^/s (standard deviation: 0.49 µm^2^/s, figure 3C). In fact, D_eff_ in the cytosol after 120 min was comparable to that in stress granules during early stress. Thus, the decrease in mobility in the cytoplasm and nucleus between 40 min and 120 min after stress induction was far more dramatic than observed in SGs. This reduction throughout the complete cytosol and nucleus was unexpected and may indicate an aggregation and oligomerization pathway independent of stress granules.

To get more detailed insight into the contribution of different TDP-43^Halo^ species (fast, medium, slow) on the mobility reduction, the different diffusion states and the respective fractions were analyzed (Fig. 3D).

Figure 3D shows, that TDP-43^Halo^ mobility reduction is caused by a decrease of the fast diffusion coefficient (D_3_) accompanied also by a reduction of the fraction of fast TDP-43^Halo^ species (F_3_) (Fig. 3D) resulting in an increase of both slow fractions, F_2_ and F_1_. Together, this suggests a general slow-down of TDP-43^Halo^ with increasing stress duration, caused by mobility reduction and the formation of less-mobile TDP-43 species.

Several control measurements were conducted. As depicted in supplementary figures S6 A and B, no reduction of the effective diffusion D_eff_ coefficient was observed for the C- and N-terminally tagged TDP-43 constructs during 120 min of measurement time under unstressed conditions. These results verify that the tracking environment or other external factors do not slow down TDP-43^Halo^. Also the mobility of the HaloTag alone was assessed under stressed conditions. In this case, no mobility reduction was seen with increasing stress duration (Fig. S6C), thus excluding an unspecific stress related mobility reduction.

Moreover, we also observed a similar effect of TDP-43 mobility reduction for the N-terminally tagged TDP-43 construct (Supplementary fig. S4) ensuring that the observed decrease in TDP-43 mobility is independent of the HaloTag position.

Table 1 gives an overview of the effective diffusion coefficient D_eff_ of the ^Halo^TDP-43 and TDP-43^Halo^ constructs at different time-points (unstressed, 60 and 120 min of sodium-arsenite stress, whole cell). The comparison shows, that ^Halo^TDP-43 generally displays a lower mobility than the TDP-43^Halo^ construct (as expected due to the lack of fragments in the case of ^Halo^TDP-43) and that also for ^Halo^TDP-43 a stress-related slow-down of mobility is observed.

**Table 1:**
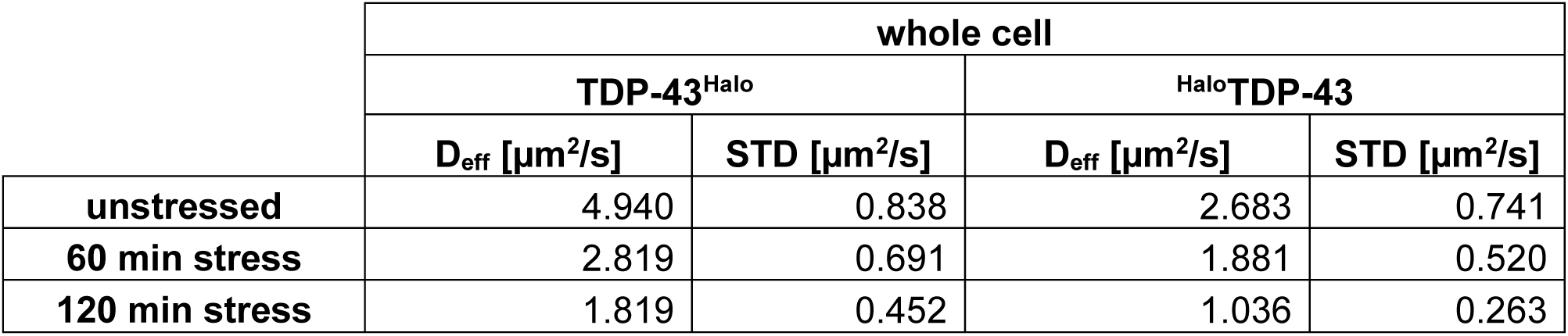
Comparison of the effective diffusion coefficient D_eff_ between TDP-43^Halo^ and ^Halo^TDP-43 constructs at different stress durations (unstressed, 60 min and 120 min 0.5 mM incubation with sodium arsenite, error given as the standard deviation (STD)). Diffusion coefficients are given for the whole cell.

|  | whole cell |  |  |  |
| --- | --- | --- | --- | --- |
|  | TDP-43<sup>Halo</sup> |  | HaloTDP-43 |  |
| | $D_{\text{eff}}$ [$\mu\text{m}^2/\text{s}$] | STD [$\mu\text{m}^2/\text{s}$] | $D_{\text{eff}}$ [$\mu\text{m}^2/\text{s}$] | STD [$\mu\text{m}^2/\text{s}$] |
| **unstressed** | 4.940 | 0.838 | 2.683 | 0.741 |
| **60 min stress** | 2.819 | 0.691 | 1.881 | 0.520 |
| **120 min stress** | 1.819 | 0.452 | 1.036 | 0.263 |

Taken together, we found region-specific effects of stress induced TDP-43^Halo^ slow down within different cellular regions. Within stress granules TDP-43^Halo^ shows, already for short stress durations, an expected, strong reduction of mobility as compared to TDP-43^Halo^ in the cytoplasm under unstressed conditions, accompanied by a further, moderate decrease in mobility with prolonged stress. Surprisingly, we also found pronounced decreased TDP-43 mobility in the nucleus and the cytoplasm upon sodium arsenite stress, suggesting that aggregation or oligomerization also occurs outside of stress granules.

To investigate the role of ALS-causing TDP-43 mutations on TDP-43 mobility, we engineered one mutation in the alpha-helical structure (M337V, (Feneberg et al. 2020; François-Moutal et al. 2019)) and another mutation in the Glycine-/Serine-rich domain (A382T, (Buratti 2015; François-Moutal et al. 2019)) (Fig. S7, S8). Single-molecule tracking did not show any significant alterations in the course of mobility of mutant TDP-43 as compared to the wild-type constructs (Fig. S8, S9), suggesting that the selected mutants do not have an additional effect on TDP-43 mobility with increasing stress. Before stress application and at low stress conditions we saw a faster mobility for familial mutants M337V and A382T (Fig. S8). Together, these results suggest that the pathological effect of the A382T and M337V mutations may not be based on an overall faster aggregation of mutated TDP-43 species.

### Longer stress durations lead to less efficient recovery of slow TDP-43 species

To find out whether and to which extent the reduced mobility of TDP-43^Halo^ can be reversed, we again employed our single-molecule tracking analysis and bulk solubility assessment. Reversibility of TDP-43 slow-down was assessed by stressing TDP-43^Halo^ cells for 60 min and 120 min with sodium arsenite and subsequent single-molecule tracking until 4 h after stress removal (Fig. 4 A - F; statistical evaluation figure S5). While short stress exposure (1h) seems to allow almost complete restoration of TDP-43^Halo^ mobility after 4h of recovery, longer stress exposure (2h) initiates processes hindering complete amelioration of TDP43 mobility reduction (Fig. 4 A, B and S5 A).

**Figure 4:**
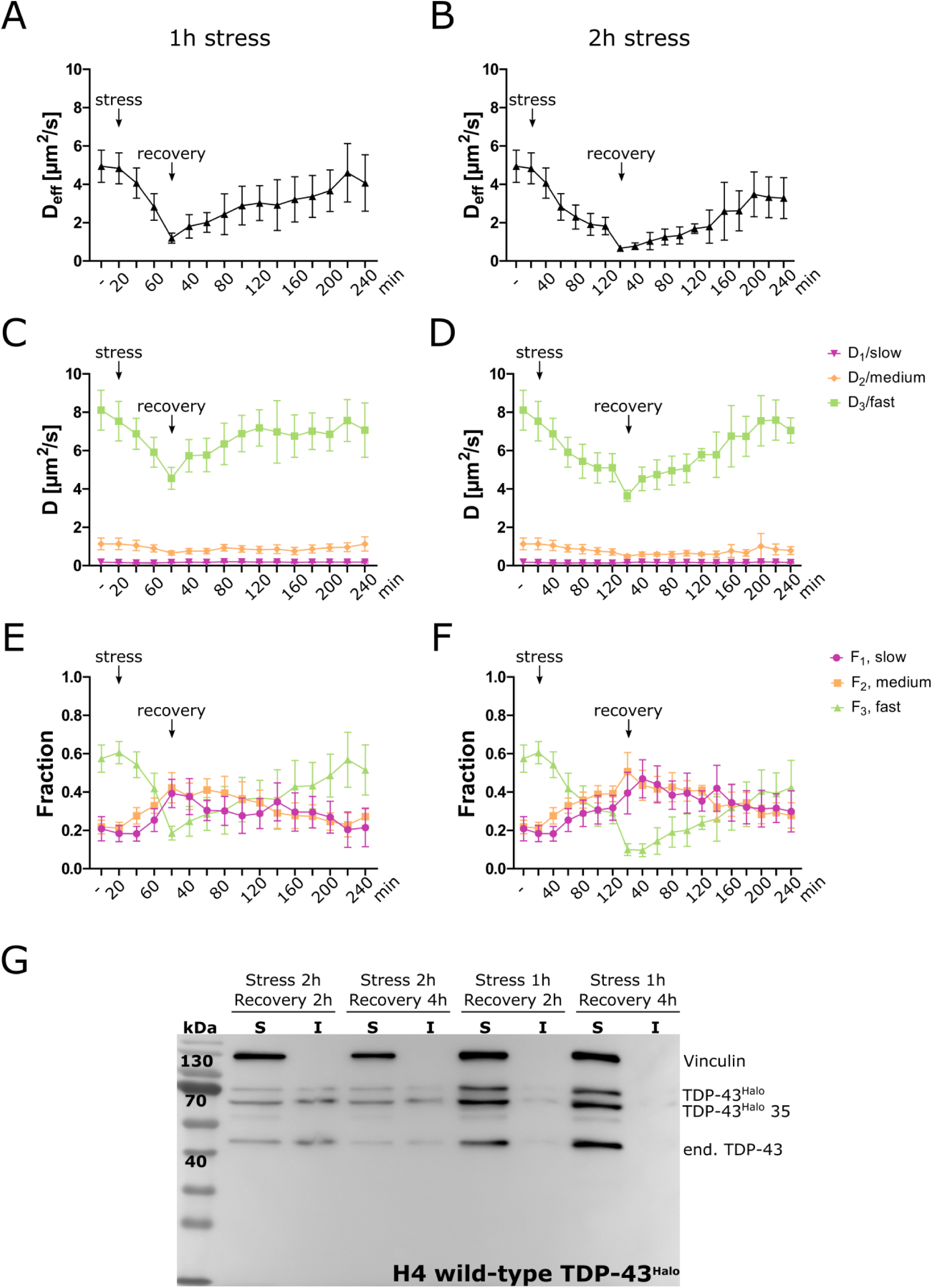
Stress and recovery experiment. A, C and E: Stress and recovery time courses of the effective diffusion coefficient D_eff_ and the diffusion coefficients D_1_ (slow), D_2_ (medium) and D_3_ (fast) and their respective amplitudes (F_1_, F_2_, F_3_) plotted for the whole cell for 1h stress duration and up to 4h of recovery (green: D_3_/F_3_, orange: D_2_/F_2_, red: D_1_/F_1_). Stress and recovery start points are marked by arrows. B, D and F: Stress and recovery time courses of the effective diffusion coefficient D_eff_ and the diffusion coefficients D_1_ (slow), D_2_ (medium) and D_3_ (fast) and their respective amplitudes (F_1_, F_2_, F_3_) plotted for the whole cell for 2h stress duration and up to 4h of recovery (green: D_3_/F_3_, orange: D_2_/F_2_, red: D_1_/F_1_). Stress and recovery start points are marked by arrows. G. Solubility assay of the TDP-43^Halo^ wild-type after different stress and recovery durations (1: Stress 2h, Recovery 4h, 2: Stress 2h, Recovery 2h, 3: Stress 1h, Recovery 4h, 4: Stress 1h, Recovery 1h). Antibodies: anti-vinculin, anti-TDP-43.

To get more insight, we again turned to the detailed analysis of the diffusion constants and respective fractions (D_1_/F_1_/slow, D_2_/F_2_/medium, D_3_/F_3_/fast) to test whether a reduced mobility is caused by a general slow-down of TDP-43^Halo^ or a shift in the respective fractions. The fast diffusion coefficient D_3_ is significantly reduced after 2h stress and both recovery conditions as compared to the unstressed condition (Fig. 5S B). For the 1h stress condition, D_3_ is significantly decreased after 120 min of recovery. After 240 min of recovery the D_3_ adapts comparable values for both stress conditions, although the decrease for the 1h stress condition was no longer significant with respect to the unstressed condition.

**Figure 5:**
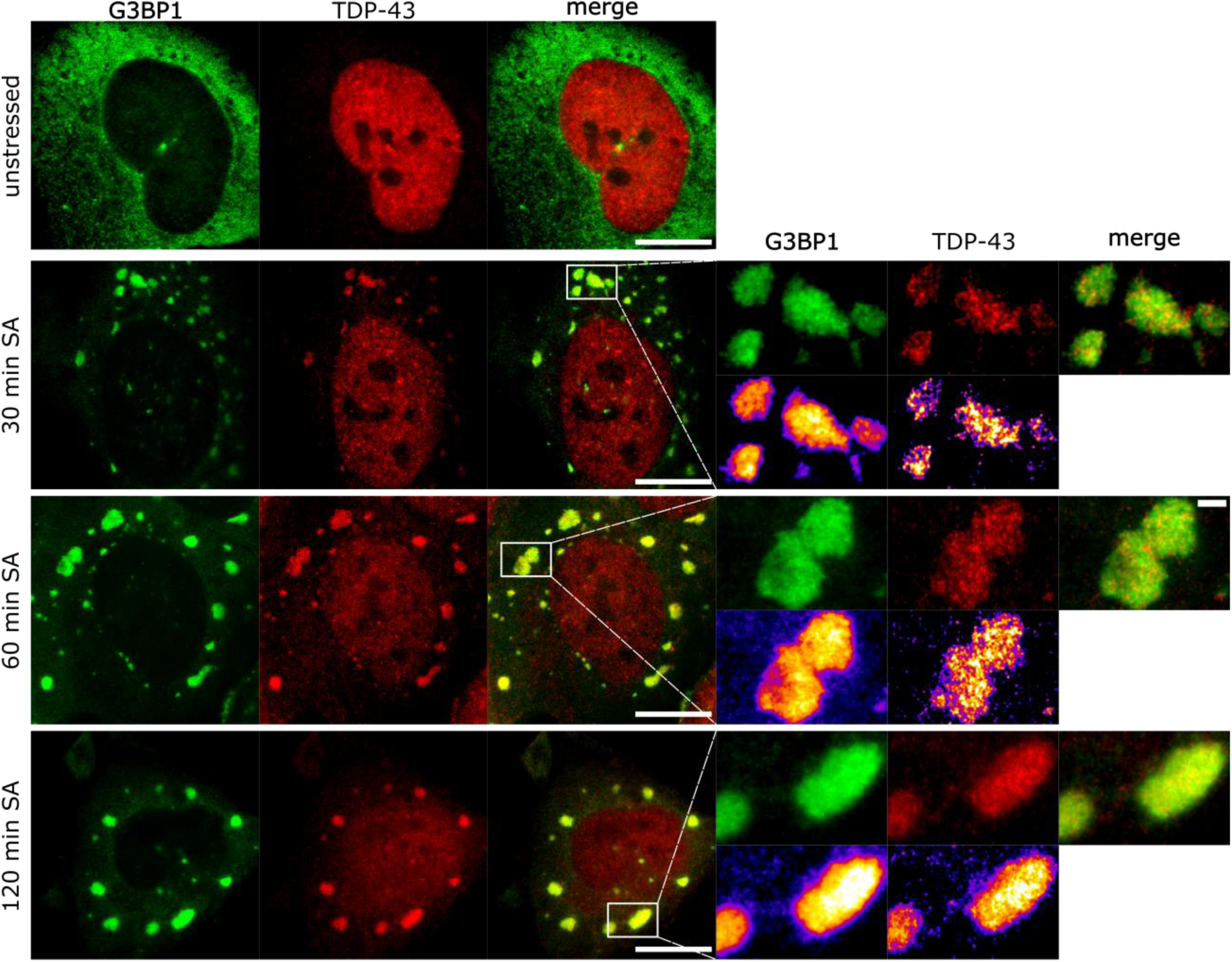
STED super-resolution microscopy reveals an inhomogeneous distribution of TDP-43 within stress granules. STED super-resolution image of unstressed H4 cells or imaged under different stress durations (30 min, 60 min, 120 min). Stress-granule crops are marked with a white rectangle (red: TDP-43-Atto647N, green: G3BP1-Atto594, scale bar 10 µm, 2 µm). For a better visualization of the substructure distribution a FIRE-LUT was used for the stress granule crops.

The fast fraction F_3_ is significantly reduced after 120 min of recovery (Figure 5S B) and longer stress duration leads to a lower fast fraction than shorter stress duration (1h stress: F_3_ = 0.37, 2h stress: F_3_ = 0.24). While 240min after stress, F_3_ shows complete recovery for cells stressed for 1h, for longer stress duration F_3_ remains reduced even after this long recovery. This again underlines the interplay between two different effects that lead to a reduced TDP-43 mobility: 1) a decrease in mobility, and 2) an increase of immobile TDP-43^Halo^ species.

To further biochemically characterize the nature of the reduced mobility after recovery, the previously described biochemical solubility assay was performed after either 60 min or 120 min of sodium arsenite stress and 2h or 4h of recovery, respectively (Fig. 4 G). After longer stress exposure, TDP-43^Halo^ shows a significant insoluble fraction, even after 4h of recovery, while after short stress almost no insoluble TDP-43 is detected (Fig. 4G).

Together, single-molecule tracking data and biochemical characterization indicate that TDP-43^Halo^ is capable to recover from short stress insults while longer stress leads to persistent, insoluble TDP43 aggregates.

### STED super-resolution imaging reveals inhomogeneous distribution of TDP-43 within stress granules

Previous studies indicated, that stress granules are not necessarily homogenous phase-separated compartments but can exhibit regions of higher density termed ‘core’ that are surrounded by a less dense ‘shell’ (Jain et al. 2016; Niewidok et al. 2018). Using super-resolution microscopy, it was reported, that stress granule components like G3BP1 or poly(A)-RNA localize in a distinct substructure within stress granules (Jain et al. 2016; Wheeler et al. 2016). Thus, we were interested if TDP-43 exhibits a similar substructure within stress granules at different stress time-points.

To obtain such high-resolution sub-compartmental spatial information immunolabelled TDP-43 and G3BP1 were imaged using stimulated emission depletion (STED) microscopy under unstressed and different stress durations (30 min, 60 min and 120 min of 0.5 mM sodium arsenite) in naïve H4 cells (Materials and Methods). TDP-43 shows an inhomogeneous distribution with denser regions (higher intensity) and less dense regions (lower intensity) within stress granules (Fig. 6). This supports the idea of an inhomogeneous distribution of TDP-43 within G3BP1-positive stress granules. In contrast G3BP1 appears more homogenously distributed throughout the stress granules. This could in part be attributed to a more saturated fluorescence signal due to the high G3BP1 density within stress granules.

**Figure 6:**
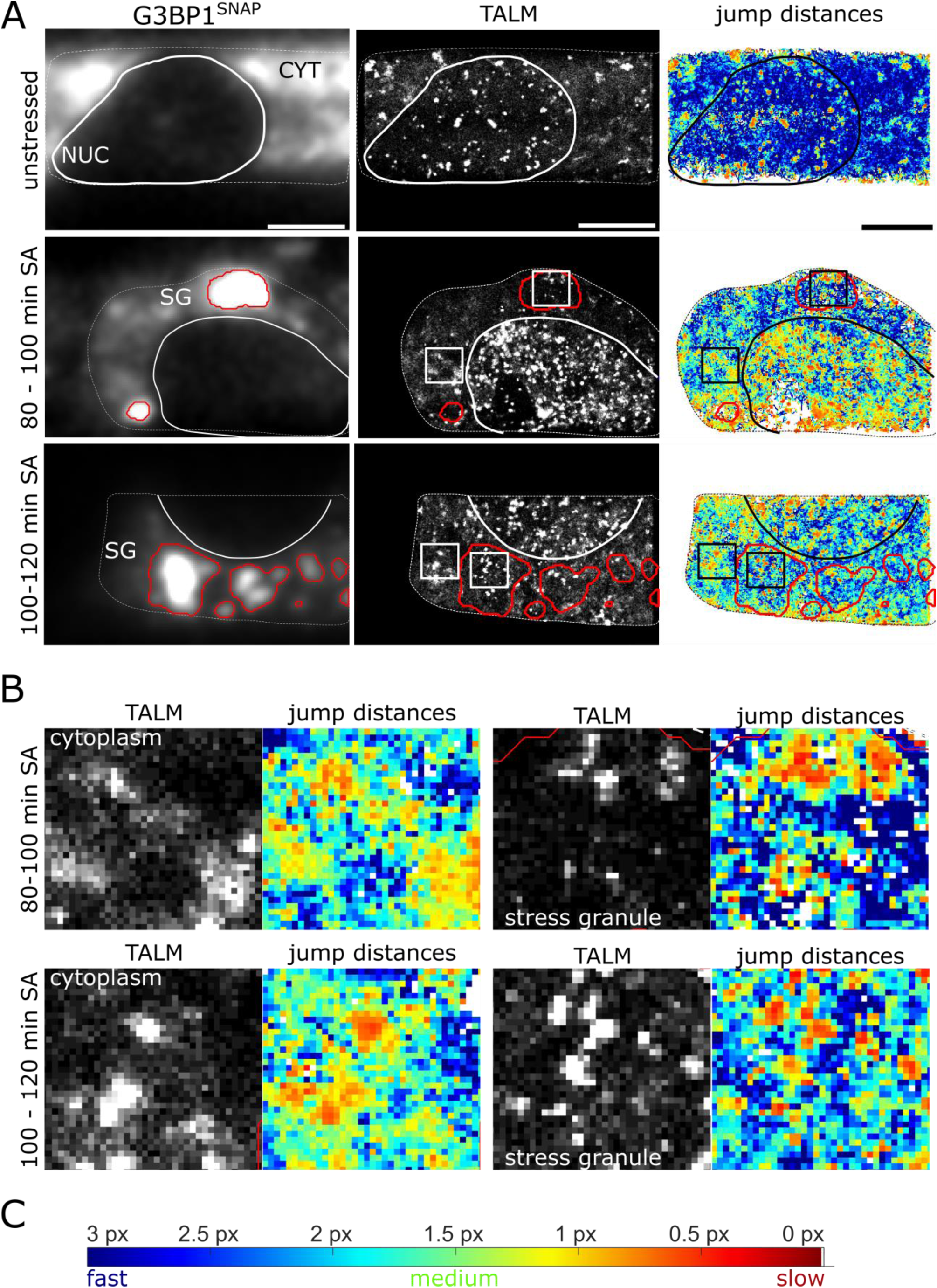
TALM (tracking and localization microscopy) and displacement mapping (DM) analysis of TDP-43 movement in H4 cells. A. Overview of images obtained with TALM and DM analysis for TDP-43^Halo^ imaged under unstressed and stressed (80-100 min and 100-120 min sodium arsenite treatment) conditions (color map: blue = fast movement, red = slow movement). B. Crops of TALM and DM images of regions within the cytoplasm and stress granules. Local binding hotspots (TALM, white spots) correlate with a reduced TDP-43 mobility (DM, red regions) in both regions. C. Color bar for the DM analysis. Blue color depicts faster movement and red color depicts slow movement. The maximum allowed jump distance for the analysis was 5 pixels. For better visualization, all jumps greater than 3 pixels was displayed in dark blue.

The highest level if inhomogeneity of TDP-43 within stress granules was visible after 30 min and 60 min of sodium-arsenite stress. After 120 min of stress the distribution of TDP-43 is more homogenous, similar to the one of G3BP1. Stress granules are known to coalesce over the stress period (Wheeler et al. 2016). A densification and increased amount of TDP-43 within stress granules could be the reason for the observed loss in substructure with increased stress.

### Single-molecule analysis shows TDP-43 oligomerization independent of stress granules

Single molecule tracking analysis showed a slow-down of TDP-43^Halo^ movement with increasing stress duration. A reduced mobility can be caused by several processes e.g. by localization to a confined compartment, by interaction with other protein complexes, by pathological aggregation or by physiological interactions. Also solubility assessment of TDP-43 under different stress condition (Fig. 3 B), confirmed an insoluble fraction with increasing stress exposure. To study the spatial distribution of the different TDP-43 mobility regimes, we performed TALM analysis (tracking and localization microscopy (Niewidok et al. 2018; Appelhans et al. 2012; Manley et al. 2008)) and investigated local jump distances (distance mapping, DM) as a means to obtain a super-resolved diffusivity map (Kuhn et al. 2021; Xiang et al. 2020).

In TALM analysis, the fitted position of every detected spot is marked and thereby a super-resolved image can be created from the tracking data. More frequent localization of TDP-43 at a given location, indicates binding to cellular structures or aggregation events. Frame to frame position jumps of localized molecules result in average displacements for each pixel in the image (Kuhn et al. 2021), indicating regions of local high or low mobility. The combination of these two methods allows for a correlation between binding hotspots or regions with increased TDP-43 localization and local mobility patterns. The visualization of such patterns can help to elucidate the origin of the observed TDP-43 slow-down in the cytoplasm.

Figure 6A shows the G3BP1 image and the respective TALM and DM image for unstressed and stressed conditions. The G3BP1 image was used to assign the cellular regions, in particular the location of stress granules (see Materials and Methods). To clearly separate stress granules from the stress-granule-free cytoplasm, a conservative approach in stress granule detection was chosen, resulting in a slightly overestimated stress granule area (see Materials and Methods). For the displacement mapping (DM) images, all jumps greater than 3 pixels (390 nm) are depicted in dark blue, thus, higher mobility correlates with a blue color and lower mobility correlates with a red color scheme (Fig. 5C).

Under unstressed conditions, TDP-43^Halo^ shows numerous binding hotspots in the nucleus and stress granules and regions of increased TDP-43 localization, termed localization patches, in the cytoplasm, that all correlate with local low TDP-43^Halo^ mobility (red).

Binding hotspots and localization patches appear as bright spots or regions within TALM images, originating from frequent TDP-43 localizations from one but typical several TDP-43 molecules. Under unstressed conditions, displacement mapping clearly shows that despite the binding hotspots and localization patches, TDP-43 diffusion is strongly dominated by high TDP-43^Halo^ mobility (blue dominated DM image) (Fig. 6A, upper panel). Such a mobility pattern of TDP-43 could be explained by the physiological shuttling between the nucleus and cytoplasm (Ayala et al. 2008).

For longer stress duration (80 - 100 min, 100 - 120 min), DM shows a shift towards medium and low TDP-43^Halo^ displacements throughout the whole cell. This fits well to the observed TDP-43^Halo^ mobility reduction in all cellular compartments obtained with single-molecule tracking. To get more detailed insight into the TDP-43^Halo^ behavior in different regions, figure 6B shows exemplary cropped areas from the cytoplasm and stress granules. TDP-43^Halo^ shows distinct binding hotspots within stress granules under both stress conditions. These observations can be explained by localized ‘binding regions’ within stress granules as previously observed for G3BP1 and IMP1 (Niewidok et al. 2018) and are consistent with the inhomogeneous stress granule structure observed with STED microscopy (Fig. 5). The binding hotspots in stress granules correlate with a low TDP-43^Halo^ mobility in DM (red spots, Fig. 6B). These binding hotspots are caused by the repeated localization of TDP-43 to these regions since an overlay between a TALM image and track start points from all molecules localized in one movie shows that several TDP-43^Halo^ tracks are originating from binding spots detected within stress granules (Supplementary figure S9). The same is observed for the cytoplasm, however, here a more dispersed distribution of track start points can be seen. Additionally, supplementary figure S9 B depicts a kymograph of a stress granule region, showing repeated binding of several TDP-43^Halo^ molecules during the measurement period.

Based on the unexpected finding that also in the cytoplasm reduced mobility of TDP-43^Halo^ was observed with increasing stress, the spatial distribution of TDP-43^Halo^ in the cytoplasm was of special interest. Figure 6B shows patches of frequent TDP-43^Halo^ localization within the cytoplasm (TALM) that correlate with a reduced TDP-43^Halo^ mobility (DM). Most importantly, throughout the whole cytoplasm the mobility is reduced. These data can be explained by a stress granule independent oligomerization mechanism distributed widely across the cytoplasm.

Additionally, TALM and DM analysis show frequent TDP-43 localization accompanied by a reduced TDP-43^Halo^ mobility to a vesicle-like compartment (example shown in Fig. 7). Interestingly, TDP-43 is not equally distributed throughout the whole vesicle, but is localized more often at the outside than at the center of the vesicle (Fig. 7A). Single TDP-43^Halo^ tracks also show TDP-43 movement along the vesicle outline, seemingly avoiding the interior (Fig 7B).

**Figure 7:**
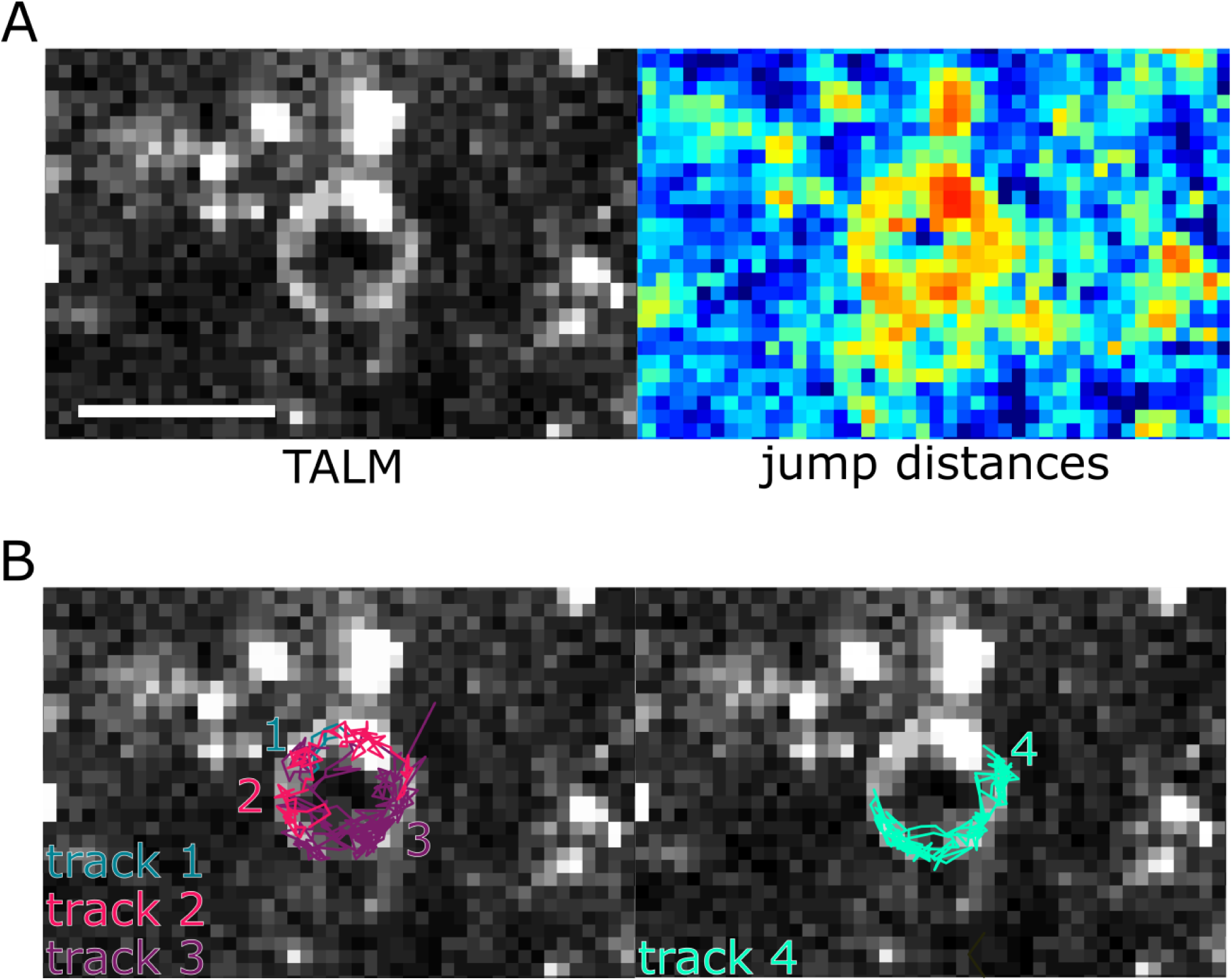
Tracking and localization microscopy (TALM) and displacement mapping (DM) analysis of vesicular structures. A. TALM and DM images of a vesicular structure found in the cytoplasm. B. Selected TDP-43^Halo^ tracks show confined movement along the outline of the vesicle depicted in A. Scale bar 1 µm.

Taken together, TALM and DM analysis are valuable tools for the visualization and better interpretation of single-molecule tracking data. With these tools, we were able to show patches of increased TDP-43^Halo^ localization in the cytoplasm that additionally correlate with a decreased jump distance, indicative of a reduced mobility. We could demonstrate that TDP-43^Halo^ is capable to oligomerize in the cytoplasm independent of stress granules.

## Discussion

The present study provides direct evidence that TDP-43 is capable to aggregate or oligomerize in the cytoplasm independent of stress granules. Using live cell single molecule tracking as well as spatial analysis of single molecule tracking data we found numerous patches with reduced TDP-43 mobility in the cytoplasm, which we interpret as sites of oligomerization.

Our newly generated stable, double-transgenic human H4 neuroglioma cell lines expressing Halo-tagged TDP-43 and SNAP-tagged G3BP1 serve as a model system that allow studying stress related changes of TDP-43 mobility and localization on a single-molecule level. To our surprise, stress induced TDP-43 mobility reduction was not limited to stress granules, but instead mobility in the nucleus as well as the cytoplasm were strongly reduced with increasing stress duration.

The mobility reduction of TDP-43 observed within stress granules was expected due to the intrinsically high viscosity and high valency of interaction partners within SGs. However, TDP-43 did only show a moderate further decrease in mobility during prolonged. Since distinct binding hotspots were already shown for other stress granule proteins like G3BP1 and IMP1 (Niewidok et al. 2018) we wanted to investigate the sub-structural distribution of TDP-43 within stress granules. The application of tracking and localization microscopy (TALM) and displacement mapping (DM) analysis showed binding hotspots that correlate with reduced mobility and kymograph analysis further highlights that these hotspots were repeatedly visited by several TDP-43^Halo^ molecules. These results further support the idea that stress granules exhibit distinct ‘core’ or binding regions (Niewidok et al. 2018; Jain et al. 2016). In addition, STED super-resolution microscopy showed that TDP-43 exhibits sub-structure within stress granules and becomes more homogenously distributed with increasing stress duration, which might be attributed to an increasing concentration of TDP-43 within stress granules at longer stress periods. A changing sub-structural distribution of TDP-43 and interaction with stress granules could elucidate the role of stress granules in the pathway of TDP-43 aggregation. Taken together, TDP-43 shows a sub-structural distribution within stress granules that might reflect possible binding sites observed using TALM imaging.

Next, we showed that the reduction of TDP-43 mobility is reversible after short stress duration. Longer stress durations lead to a reduced TDP-43 mobility after recovery, that can be attributed to a reduction in the diffusion coefficient and an increase of slower TDP-43 species. This finding is paralleled by the formation of insoluble TDP-43 species at longer stress durations seen by biochemical fractionation, suggesting that reversibility of TDP-43 aggregation and insolubility strongly depends on the stress duration applied. The impairment of protein degradation systems like autophagy or the ubiquitin proteasome system, may lead to TDP-43 aggregation or a defective aggregate clearance (Scotter et al. 2014). Whether impaired protein degradation is causative for the reduced recovery capability of TDP-43^Halo^ was not part of this study and has to be investigated in future experiments.

Although a reduced mobility of TDP-43 within stress granules was expected, the strong reduction of TDP-43 mobility in the nucleus and especially the cytoplasm was surprising. 120 min of sodium arsenite stress, lead to a reduction of TDP-43 mobility of about 50 % in the cytoplasm. Assuming no changes in the effective viscosity of the surrounding medium, such a decrease in diffusivity would correspond to a doubling of the hydrodynamic radius and thus to an 8-fold increase of the effective mass and volume of the protein complex. These results might therefore suggest the formation of TDP-43 oligomers under the observed conditions. Importantly, oligomer formation is not localized to stress granules, but occurs throughout the nucleus and the cytoplasm. These results are also in line with our previous work demonstrating TDP-43 oligomerization after sodium arsenite stress using a TDP43 protein complementation assay, leading to the formation of high molecular weight oligomers (Feiler et al. 2015). This results might be explained by the mode of action of sodium arsenite leading to reduced cellular ATP levels and mitochondrial impairments (Yih et al. 1991; Shi, Shi, and Liu 2004; Jomova et al. 2011). Moreover, recently also cytoplasmic TDP-43 aggregates and the formation of insoluble TDP-43 was shown after oxidative stress (Zuo et al. 2021). These TDP-43 aggregates additionally sequester mitochondrial proteins, leading to mitochondrial dysregulation, further promoting oxidative stress and aggregation (Zuo et al. 2021). These data nicely support the data presented in this study, attributing an important role to inhibition of glycolysis in TDP-43 cytoplasmic aggregation and provide supporting evidence of the formation of SG independent TDP-43 aggregates.

In line with these observations are the tracking and localization microscopy (TALM) and displacement mapping (DM) analysis data, showing local patches of reduced TDP-43 mobility in the cytoplasm correlating with increased TDP-43 localization seen by TALM microscopy. It is interesting to mention that the spatial distribution of TDP-43 within the cytoplasm strongly differs from that observed in stress granules. Within stress granules, distinct binding hotspots were observed, whereas within the cytoplasm, TDP-43 shows an almost homogeneous slow-down of TDP-43 molecules with some more pronounced localization patches. We therefore hypothesize that the oligomerization observed throughout the cytoplasm might be a first step during aggregate formation.

Our data on TDP43 mobility reduction in the cytoplasm are in line with two recent studies demonstrating in other model systems TDP-43 aggregation or oligomerization in the cytoplasm (Mann et al. 2019), (Gasset-Rosa et al. 2019). Using an optogenetic clustering system to induce controlled phase separation of TDP-43 wild-type and RNA binding deficient mutants, (Mann et al. 2019) report, that exclusively cytoplasmic, RNA-binding deficient TDP-43 forms stress granule independent aggregates, that exhibit pathological features like hyper-phosphorylation and are devoid of RNA. TDP-43 localized to these inclusions did exhibit a reduced recovery capability in FRAP measurements as compared to TDP-43 within stress granules, indicating aggregation.

(Gasset-Rosa et al. 2019) employed an exclusively cytoplasmic Δ-NLS-TDP-43 construct and investigated TDP-43 phase separation after the application of fragmented TDP-43 or FUS particles as well as additional sodium arsenite stress. They showed, that exogenously applied TDP-43 of FUS particles can induce aggregation of endogenous TDP-43 and that these aggregates mostly, do not co-localize with stress granule markers. Additionally, they were able to show that after sodium arsenite treatment, Δ-NLS-TDP-43 first localizes to stress granules but forms stress granule independent assemblies at later time points. Similar to what was seen in (Mann et al. 2019), these assemblies did exhibit a reduced recovery in FRAP experiments as comparted to stress granule localized TDP-43.

Although both studies used different stress models and TDP-43 constructs, they could independently show TDP-43 phase separation and aggregation in the cytoplasm, highlighting the importance of cytoplasmic TDP-43 oligomerization in ALS pathogenesis and research. The results of both studies, (Mann et al. 2019; Gasset-Rosa et al. 2019), support the results obtained in this study. There are additional, important surveys that showed that TDP-43 aggregation can occur independently of stress granule formation or localization (Fernandes et al. 2020; Hans, Glasebach, and Kahle 2020). Lastly, (McGurk et al. 2018) showed that TDP-43 stress granule localization is mediated by PAR-binding and that PAR-binding deficient mutants are no longer able to translocate to stress granules and subsequently form aggregates. All these studies attribute a rather protective role to TDP-43 stress granule localization and RNA interactions and highlight the importance of studying TDP-43 also outside of these compartments. This effect might not only be applicable for TDP-43, since (Shelkovnikova et al. 2013) showed a similar, protective effect of stress granule localization for the RNA-binding protein FUS.

ALS pathology was shown to spread in patient’s brains (Braak et al. 2013) and the search for a pathological, spreading species is a highly relevant topic. Previously we have shown evidence for cellular TDP-43 oligomerization by complementation essays and size exclusion chromatography (Feiler et al. 2015) and we speculated that TDP-43 oligomers might serve as pathogenic species for propagation of pathology. In addition, TALM analysis revealed that TDP-43 can localize to confined, vesicle-like structure, where TDP-43^Halo^ was localized exclusively to the outer layer of this structure. Similar TDP-43 containing structures were already seen for nuclear TDP-43 (Yu et al. 2021; Wang et al. 2020).

The reduction of D_eff_ of TDP-43 in the nucleus could be explained by an increased nucleic acid binding with increasing stress duration or a shift of the mobile TDP-43 fraction to the cytoplasm, which is a commonly observed feature of TDP-43 under stressed conditions (Ayala et al. 2008; Correia et al. 2015; Giordana et al. 2010). Tracking and localization microscopy (TALM) showed, that under stressed conditions, TDP-43^Halo^ exhibits distinct and frequent binding hotspots within the nucleus. This argues for an increase in TDP-43 binding or aggregation within the nucleus with longer stress, which is in accordance with several studies reporting on TDP-43 phase-separation in the nucleus under unstressed and stressed conditions (Gasset-Rosa et al. 2019; Yu et al. 2021; Wang et al. 2020) and our data gives further quantitative evidence for these effects.

In conclusion, we were not only able to demonstrate a stress related slow-down of TDP-43 mobility within stress granules, but also a reduction in TDP43 mobility in the cytoplasm independent of stress granules and in the nucleus. TDP-43 shows a stress related and reversible decrease of mobility in all cellular compartments, indicating stress related binding and oligomerization events. The unexpected prominent and inhomogeneous reduction of TDP-43 mobility in patches of high TDP-43 concentration in the cytoplasm strongly argues for a probably aggregation promoting dynamic of TDP-43 protein distinct from TDP-43 located in SGs.

## Material and Methods

### Cloning procedure

The phage UbiC G3BP1-SNAP plasmid was purchased from Addgene (Plasmid #119949) and characterized and published previously (Wilbertz et al. 2019). The TDP-43 constructs or the HaloTag only construct were cloned into an LVTetO lentiviral backbone (Gebhardt et al. 2013). An insert was cloned into the plasmid to introduce necessary restriction sites. The used primer and insert sequence are listed in Table 1. To generate the TDP-43 mutants M337V and A382T, single point mutations were introduced to the respective wild-type plasmids by site-directed mutagenesis using the Q5 site-directed mutagenesis Kit (NEB). The used primers are listed in table 1.

**Table 1:**
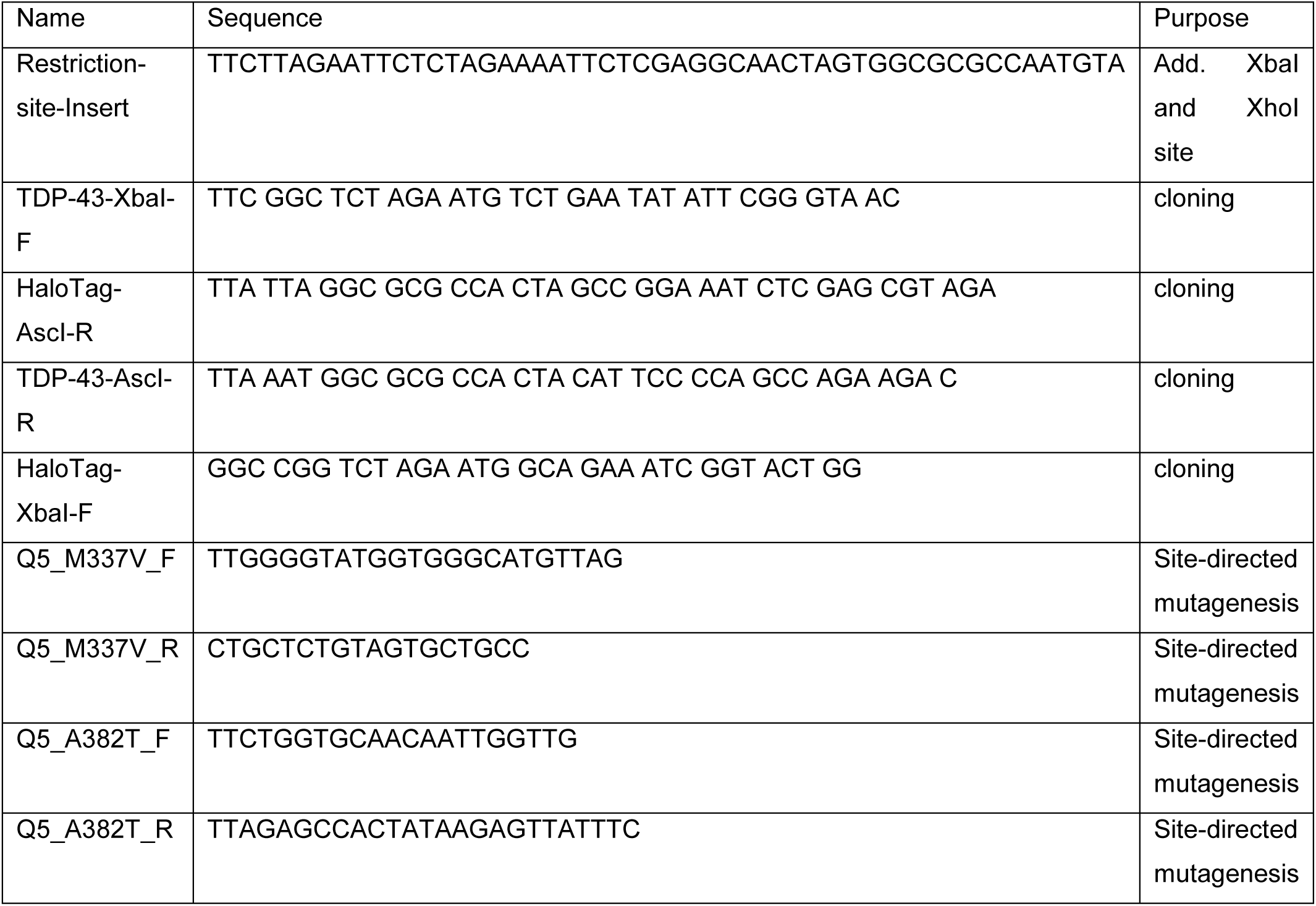
Used Primer

### Stable cell line generation

LentiX cells (subclone of the HEK293T cell line), necessary for lentiviral particle production, were thawed one week before transfection. LentiX cells were grown to 80 % confluence on a 10 cm dish (Sarstedt) prior to transfection. Cells were transfected using 10 µg transfer plasmid containing the respective TDP-43 construct, 7.5 µg packaging plasmid psPAX2 (Addgene plasmid #12260) and 2.5 µg envelope plasmid pMD2.G (Addgene plasmid #12259) using the jetPRIME transfection system (Polyplus transfection). All three plasmids were mixed with 500 µl jetPRIME buffer and vortexed for 20 s. 30 µl jetPRIME transfection reagent was added and the mixture was again vortexed for 20 s. The mixed solution was incubated for 10 min at room-temperature and added dropwise to the LentiX cells. The transfected cells were incubated for 2 days at 37°C and 5% CO_2_ to ensure proper viral particle production.

One day before viral transduction the target H4 cells (ATCC HTB-148) were seeded on a 6-well dish to reach 60% confluence the next day. At the day of viral transduction, the supernatant (9-10 ml) containing the viral particles was removed and 500 µl of the supernatant was added to the H4 cells. For the generation of the double-transgenic cell lines 500 µl of viral particles containing the G3BP1-SNAP construct and 500 µl containing the respective TDP-43-Halo construct were added to one well respectively. To reach proper transduction efficiency the wells were incubated for 72h at 37°C and 5% CO2. After 72 h the cells were once washed with DPBS, trypsinized and transferred onto a 10 cm dish. To remove cells without or with only one construct, the cells were sorted for double-positive clones at the Core Facility Cytometry at Ulm University.

### Cell line cultivation

Sorted cell lines were cultivated in DMEM supplemented with 10 % FBS, and 1 % sodium-pyruvate, 1 % non-essential amino-acids and 1 % Glutamax. The cell lines were kept in culture up to 10 passages. After that they were discarded and a new vial of frozen cells was taken into culture.

### Western Blot sample preparation

#### RIPA Lysates

For lysate preparation of unstressed cell lines RIPA buffer (sigma-aldrich, R0278) was used. Cells were seeded on a 10 cm dish and grown to 80-90 % confluence. For lysate preparation, cells were washed with PBS, trypsinized and transferred into a 15 ml falcon. The cell suspension was centrifuged for 5 min at 2000 g (Centrifuge Beckman Coulter Allegra X-15-R, Rotor Sx4750A)) and 4°C. The supernatant was removed and the pellet was washed with 5 ml ice-cold PBS. The suspension was again centrifuged for 5 min at 2000 g and 4°C. The supernatant was discarded. The pellet was dissolved in 400 µl ice-cold RIPA buffer and transferred into a 1.5 ml Eppendorf tube. The lysate was incubated for 30 min on a rotating wheel at 4°C and centrifuged at 16 000 g for 20 min at 4°C (Eppendorf centrifuge 5417-R, Rotor F45-30-11). The supernatant was transferred to a fresh 1.5 ml Eppendorf tube and the pellet was discarded. The protein concentrations were adjusted to 1 mg/ml using a BCA assay (Thermo Fischer Scientific, 23227). Lysates were shock-frozen using liquid nitrogen and stored at -80°C. Triplicates were prepared for each cell line.

For western blotting 24 µl protein lysate (1 mg/ml), 3.7 µl 1M DTT and 9.3 µl 4x LDS were mixed, heated for 5 min at 95°C and stored at -20°C until usage.

#### Solubility Assay

Cells were seeded on a 5 cm dish and grown to 80-90 % confluence. Before the assay 5 ml fresh medium was added to the dish. The cells were either kept unstressed or treated 40 min, 80 min and 120 min with 0.5 mM sodium arsenite, respectively. After the end of the stress duration the medium was removed and the cells were washed with 2 ml PBS. The PBS was removed, 200 µl RIPA buffer was added per dish and cells were scraped of using a cell scraper. Lysed cells were transferred to a fresh 1.5 ml Eppendorf tube and sonified 3 times for 3 s each. The protein concentration was determined by a Bradford assay and adjusted to 1 mg/ml for each sample using RIPA buffer. From each sample 100 µl lysate were transferred to a special tube for ultra-centrifugation (Beckman Coulter, 1.5 ml tube, 257448). The samples were centrifuged for 30 min at 100 000 g at 4°C (Centrifuge Beckman Coulter Optima XPN-80, Rotor type 70.1). The supernatant, containing the RIPA-soluble fraction, was transferred into a fresh 1.5 ml Eppendorf tube. The pellet was washed with 200 µl RIPA, shortly sonified and centrifuged for 30 min at 100 000 g at 4°C (Centrifuge Beckman Coulter Optima XPN-80, Rotor type 70.1).). The supernatant was discarded and the pellet was dissolved in 100 µl 8 M Urea (8 M Urea, 20 mM Tris, pH 8). For better dissolution the sample was again sonified 3 times for 3s.

For western blotting 13 µl protein lysate (1 mg/ml), 2 µl 1M DTT and 5 µl 4x LDS were mixed, heated for 5 min at 95°C and stored at -20°C until usage. Triplicates were prepared for each cell line.

### Western Blotting

Protein separation by size was done via SDS-PAGE. For the cell line expression overview ready-made gels (Invitrogen NuPAGE 4-12%. NP0335, NP0336) and buffers (Running buffer NP0002, Transfer Buffer NP0006-1) were used. For protein separation polyacrylamide gels with a 12 % separating and a 5 % stacking gel were prepared.

Samples were heated at 95°C for 5 minutes before loading the onto the PAA-gel (marker: PageRuler Plus Prestained protein ladder (Thermo Scientific)). PAA-gels were pre-run for 30 min at 90 V. After that the voltage was increased to 125 V for 2 ½ h (SDS-running buffer: 25 mM Tris, 192 mM Glycin, 1g/L SDS). The gel was transferred to the blot chamber (XCell II, EI9051) and blotted on a nitrocellulose membrane (Thermo Scientific, LC2008) for 3h at room-temperature (Blotting buffer: 25 mM Tris, 192 mM Glycin, 20 % Ethanol). A proper protein transfer was confirmed by PonceauS staining and the membranes were blocked for 2h with 5% skin-milk powder in 1x TBS-T (1x TBS, 0.05 % Tween-20). Membranes were incubated overnight with primary antibodies (1:100 in 5% skim-milk powder, anti-TDP-43 (rb, Proteintech 10782-2-AP), anti-G3BP1 (ms, sigma, WH0010146M1), anti-Vinculin (ms, Abcam, ab18058)) at 4°C. Blots were washed 3 times for 20 min with 1x TBS-T to remove unbound primary antibodies. Blots were incubated with secondary antibodies for 1h at room-temperature (1:10000 in 5% skim-milk powder, anti-rb-HRP-conjugate (Promgea, W401B)/anti-ms-HRP-conjugate (Promega, W402B)). After 3 times washes for 20 min with 1x TBS-T to remove unbound secondary antibodies and bands were visualized by adding ECL solution (Thermo Fischer Scientific, 32106) using the Fusion Solo7S Imager (Vilber). The Fusion software was used to analyze overexpression of the TDP-43-Halo and G3BP1-SNAP constructs.

### Spinning disk confocal microscopy

Cells were seeded on 2-well Ibidi dishes (Ibidi GmbH, 80287) and labelled with 1 µM TMR-HaloTag ligand (Promega, G8251) and 0.12 µM SiR-SNAP-Tag ligand (NEB, S9102S) for 30 min at 37°C and 5% CO_2_. Cells were washed twice with PBS to remove unbound dye and stored in DMEM for at least 30 min prior to imaging. For imaging the medium was changed to OptiMEM. Cells were imaged at a custom-built spinning disk confocal microscope with temperature control and CO_2_ supplementation. The microscope was built around an inverted microscope body (Axio Observer, Zeiss) together with a CSU10 scan head (Yokogawa). For imaging an oil immersion objective (UPlanSApo 60x/1.35 NA, Olympus) and for detection an EMCCD camera (DV-887, Andor) were used. For excitation a 532 nm laser MGL-III-532-100 mW) and a 640 nm laser (Toptica iBeam-SMART 640-S) were used. Images were taken with a pixel size of 200 nm, an exposure time of 300 ms and averaging over 8 frames. Image analysis was performed with ImageJ.

For a comparison between naïve H4 cells and the TDP-43 Halo cells were either kept unstressed or stressed for 60 min with 0.5 mM sodium arsenite. After the stress duration cells were fixed with 3.7 % EM-grade PFA (Electron Microscopy Sciences, Catalog no. 15714) for 20 min at room-temperature. Samples were blocked with 3% BSA and 0.3 % TritonX-100 overnight at 4°C. Primary antibodies (1:500, anti-TDP-43 (rb, Proteintech 12892-1-AP), anti-G3BP1 (ms, sigma, WH0010146M1),) were incubated for 2h at room-temperature. Secondary antibodies (anti-rb-Alexa647 (Invitrogen A-21245) and anti-ms-Alexa532 (Invitrogen A11002), 1:500 each) were incubated overnight at 4°C.

### Live-cell single molecule imaging

H4 cells were seeded on a 35-mm Ibidi-dish with glass bottom (Ibidi GmbH, 81158) two days before imaging. One day before imaging cells were labelled with 10 nM photoactivatable-JaneliaFluor-646-HaloTag ligand (PA-JF646-HTL) and 0.3 µM TMR-SNAP-Tag ligand (TMR-STL) for 1h at 37°C and 5% CO_2_. After 3 washing steps with PBS for 5 minutes to remove unbound dye, cells were stored in DMEM. The medium was then exchanged 3 times during the day for fresh DMEM.

Imaging was performed on a custom-built wide-field fluorescence microscope first described here (Schoen et al. 2016). A top stage incubator system with incubation chamber (H101-MCL-NANO-ZS200, Okolab s.r.l., Italy) was used to enable temperature control and CO_2_ supplementation. Measurements were conducted at 37°C and 5 % CO_2_.

Illumination and fluorophore activation was done using 640 nm (Omikron LuxX® 638-300, 200 mW), 532 nm (Cobolt Samba, 150 mW) and 405 nm (Toptica iBEAM-SMART-405-S) laser lines. For single-molecule detection an illumination power of 2 mW (640 nm) and an activation power of ∼ 1 µW (405 nm) were used. Imaging was performed in HILO-mode with an 100x oil immersion objective (Plan APO TIRF 100x NA 1.45 Oil, Nikon). Detection was performed using EMCCD AndorSolis cameras for each channel (iXon 897 Ultra and iXon 897, Andor Technology) and an effective pixel size of 130 nm. The AndorSolis software (Version 4.31.30024.0) was used to record the tracking movies.

#### Data acquisition

Single-molecule tracking measurements were performed with an exposure time of 5 ms (total frame cycle time 6.742 ms) and under continuous low 405 nm activation of the PA-JF-646-dye. Cells were either imaged under unstressed or stressed conditions (0.5 mM sodium arsenite) for two hours. Per movie 10000 continuous frames were recorded in the tracking channel (TDP-43-PA-JF-646). Every 200 frames an image in the G3BP1-TMR channel was taken. For the TALM (tracking and localization) analysis imaging was performed for 30000 frames and 405 nm activation during the frame transfer time between each frame was used. For recovery measurements the cells were stressed for 60 min or 120 min with 0.5 mM sodium-arsenite and then imaged for 0 - 120 min and 120 - 240 min after stress removal

#### Data analysis

##### Tracking and classification of tracks into regions

Spot detection, tracking and diffusion analysis was performed using TrackIt (Kuhn et al. 2021). Spot detection was performed with a threshold factor of two. Tracking was performed using the ‘Nearest neighbor’ algorithm, with a tracking radius of five pixels, a minimum track length of three, one allowed gap frame to bridge detection gaps and a minimum track length of two frames before a gap frame. Cell outlines were used as tracking regions and additional sub-regions were drawn to define three region classes – nucleus, cytoplasm and stress granules. The cell outline and nuclear outlines were drawn manually based on the signal from the G3BP1-SNAP control channel. To account for signal fluctuations, a ‘moving average’ filter of two frames was applied. For TALM and DM data, additionally a Gaussian blurring of the G3BP1 channel of two pixels was applied.

Stress granules were outlined by an intensity-based-threshold defined by the G3BP1-SNAP control channel and were assigned dynamically over the whole movie time-course, to adapt for the mobility of stress granules. To better assess region-wise TDP-43 mobility, tracks that crossed region borders, were split at the border and assigned separately to the respective region.

##### Diffusion analysis

Diffusion coefficients and fractions were computed by fitting the cumulative distribution of displacements with a three-exponential Brownian diffusion model (Kuhn et al. 2021; Mazza et al. 2012). The bin size of the jump distance histogram was set to 1 nm. To prevent an overrepresentation of immobile molecules, a maximum of 5 jumps per track were considered while jumps over gap frames were discarded. A three component Brownian diffusion model was chosen based on the fit quality assessed by SSE and adjusted R²-error (see figure S3). Three-exponential fitting results in three diffusion coefficients D_i_ and respective fractions F_i_ -slow (D_1_/F_1_), medium (D_2_/F_2_) and fast (D_3_/F_3_). From these the effective diffusion coefficient D_eff_ was calculated:

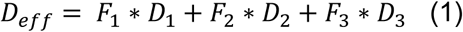

Movies where the number of jumps was less than 30 were discarded. As an additional filtering step we used the 95% confidence interval of the fit to account for erroneous fitting. Movies were discarded if the error was 5 times higher than the median error calculated from all movies within a region and a specific stress condition. Mean values and standard deviations of the diffusion coefficients, fractions and effective diffusion coefficients were then calculated from the movie-wise values of the diffusion constant or fractions of the remaining movies.

##### TALM and displacement mapping

For the visualization of the TALM and DM data a scaling factor of two was applied, increasing the number of pixels by a factor of 4 and resulting in an effective pixel size of 65 nm. In TALM analysis, the fitted position of every detected spot is assigned to the pixel in which it appeared and the detection count of the respective pixel increased by one, creating a super-resolved image from the tracking data (Manley et al. 2008). Higher count numbers (bright spots) in the TALM image indicate a more frequent localization of TDP-43 at the given spot, indicating binding to cellular structures or aggregation events. The DM analysis gives a pixel-wise average of displacement in the image (Kuhn et al. 2021), indicating local pattern with either high or low mobility (red: slow, blue: fast). All jumps greater than 3 pixels (390 nm) are depicted in dark blue. Stress granule outlines are slightly overestimated, to avoid the inclusion of signal, originating from TDP-43 within stress granules, to the cytoplasm.

### STED super-resolution microscopy

Cells were seeded on 8-well Ibidi plates with glass-bottom (Ibidi GmbH, 80827) two days before fixation. Cells were either kept unstressed or stressed using 0.5 mM sodium arsenite. After the respective stress duration (30 min, 60 min or 90 min), cells were fixed using 3.7 % EM-grade PFA for 20 min at room-temperature and washed afterwards 3 times with PBS for 5 mins. Cells were permeabilized and blocked for 2h at room-temperature (Blocking buffer: 3% BSA and 0.3 % Triton-X-100 in PBS). After primary antibodies incubation overnight at 4°C in 1:10 blocking buffer (anti-TDP-43 (rb, 1:500, Proteintech 12892-1-AP), anti-G3BP1 (ms, sigma, WH0010146M1, 1:1500,)), cells were washed 3 times for 10 min with PBS to remove unbound primary antibodies. Secondary antibodies were incubated for 1h at room-temperature (anti-rb/ms-Atto647N (sigma 40839-1ml-F/sigma 50185-1ml-F), anti-rb/ms-Atto594 (sigma 77671-1ml-F/sigma 76085-1ml-F), either 1:500 or 1:1500). Cells were then washed 3 times for 10 min with PBS to remove unbound secondary antibodies. Samples were stored at 4°C until imaging. Just prior to imaging, cells were covered in 97 % TDE.

STED images were taken using a custom-built dual-color three-dimensional STED microscope described in (Osseforth et al. 2014). For improved cross-talk reduction the emission filter in front of the 590 nm APD was changed to 623/24 BrightLine HC (AHF).

For sample illumination a randomly polarized super-continuum laser source (repetition rate 1 MHz) was split into the excitation wavelengths of 568 nm and 633 nm and their respective depletion wavelengths of 710 nm and 750 nm. A typical excitation beam power of ∼ 0.8 µw and a depletion beam power of ∼ 1.3 mW were used during image acquisition. Confocal images were taken with a pixel size of 50 nm, at a dwell time of 200 µs per pixel and peak photon number of 160 – 200 counts. STED images were taken with a pixel size of 20 nm, at a dwell time of 300 µs per pixel and a peak photon number of ∼ 150 counts. Image analysis was done using ImageJ. For better visualization Gaussian blur with sigma = 1.5 was applied.

### Statistical information

For statistical comparison of the tracking control data sets a Brown-Forsythe and Welch ANOVA test was used. For the comparison of stress time-points in the single molecule tracking data a multiple, unpaired t-test with Welch correction was used. All statistical analysis was performed using GraphPad prism version 9.0.1.

## Acknowledgements

We thank the Core Facility FACS of Ulm University for their help with cell sorting, with special thanks to Dr. Simona Ursu and Dr. Sarah Warth. The photoactivatable Janelia Fluor 646 HaloTag ligand (PA-JF 646-HTL) was kindly provided by Luke Lavis (Janelia Research Campus, Howard Hughes Medical Institute, USA). Halo-SiR ligand was kindly provided by K. Johnsson (Max Planck Institute for Medical Research, Heidelberg, Germany). We thank Ramona Bück (Neurology, University Clinic, Ulm) for her support in the cell culture and biolaboratory We thank Astrid Bellan-Koch for her help in cloning the TDP-43 mutant plasmids.

## Author contributions

JM, KMD and LS conceived the study; JM, KMD and LS designed experimental approaches; LS designed and performed cloning, cell line generation and imaging experiments; TV optical setup work and maintenance; TK derived the tracking algorithm; LS and TK, JCMG performed data analysis; JM, KMD and LS wrote the manuscript; JHW, ACL, JCMG provided intellectual input.

## Data availability

Data supporting the findings of this manuscript will be available from the corresponding author after publication upon reasonable request. All raw single particle tracking data is available in a TrackIt compatible Matlab file format on the Dryad Digital Repository. Provisional access link https://datadryad.org/stash/share/WMl6kJQhaXGefNk-JqUp6xiK3w7uGIPbLckAo4anM4Q.

The TrackIt algorithm is freely available. A Matlab version of TrackIt is available at https://gitlab.com/GebhardtLab/TrackIt/

## Material and correspondence

LS, JM, KMD

