## Supplementary information for "Stress induced TDP-43 mobility loss independent of stress granules"

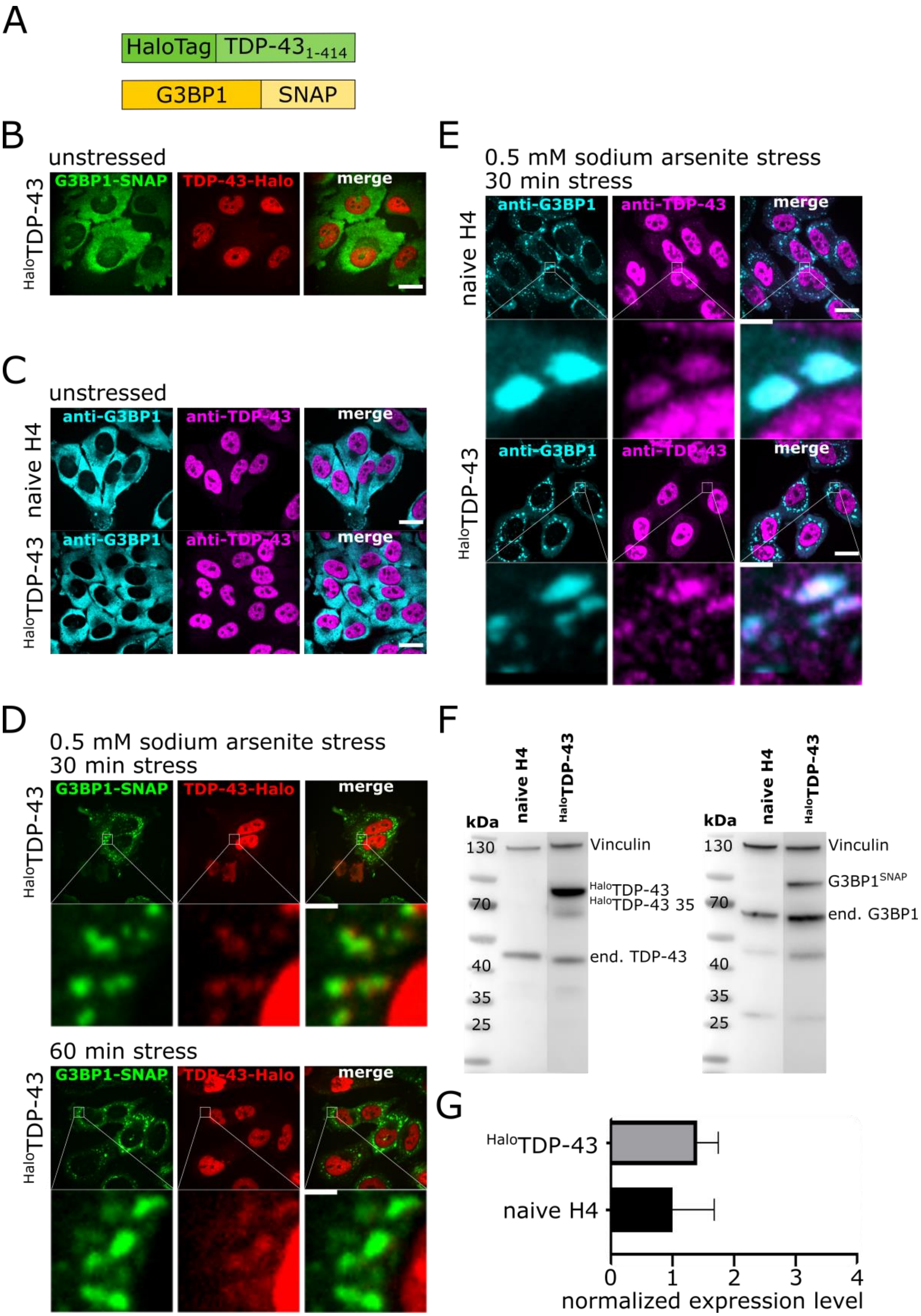
A
HaloTag
TDP-43 1-414
G3BP1
SNAP
B
unstressed
HaloTDP-43
G3BP1-SNAP
TDP-43-Halo
merge
C
unstressed
naive H4
HaloTDP-43
anti-G3BP1
anti-TDP-43
merge
D
0.5 mM sodium arsenite stress
30 min stress
60 min stress
E
0.5 mM sodium arsenite stress
30 min stress
F
kDa
naive H4
HaloTDP-43
130
70
40
35
25
Vinculin
HaloTDP-43
HaloTDP-43 35
end. TDP-43
G3BP1SNAP
end. G3BP1
G
normalized expression level
0
1
2
3
4

2

*Figure S1: Generation of a $^{Halo}$TDP-43 wild-type cell lines. A. Schematic overview of the $^{Halo}$TDP-43 construct. B. Spinning disk confocal images of the $^{Halo}$TDP-43 cell line under unstressed conditions (red: TDP-43-TMR, green: G3BP-SiR scale bar 20 µm). C. Spinning disk confocal images of naïve H4 cells under unstressed conditions (cyan: anti-G3BP1-Alexa532, magenta: anti-TDP-43-Alexa647, scale bar 20 µm). D. Spinning disk confocal images of the $^{Halo}$TDP-43 cell line under 30 min and 60 min sodium arsenite treatment (red: TDP-43-TMR, green: G3BP-SiR, scale bar 20 µm and 2 µm). E. Spinning disk confocal images of the immunostained $^{Halo}$TDP-43 cell line and naïve H4 cells under 60 min sodium arsenite treatment (magenta: anti-TDP-43-Alexa647, cyan: anti-G3BP-Alexa532, scale bar 20 µm and 2 µm). F. Western Blot overview of the $^{Halo}$TDP-43 cell line and naïve H4 cells stained with anti-vinculin, anti-TDP-43 or anti-G3BP1 antibodies showing proper expression of the transgenic constructs. G. Quantification of the overexpression for the $^{Halo}$TDP-43 cell lines compared to endogenous TDP-43 in naïve H4 cells shows a 1.5 – 2x overexpression of $^{Halo}$TDP-43.*

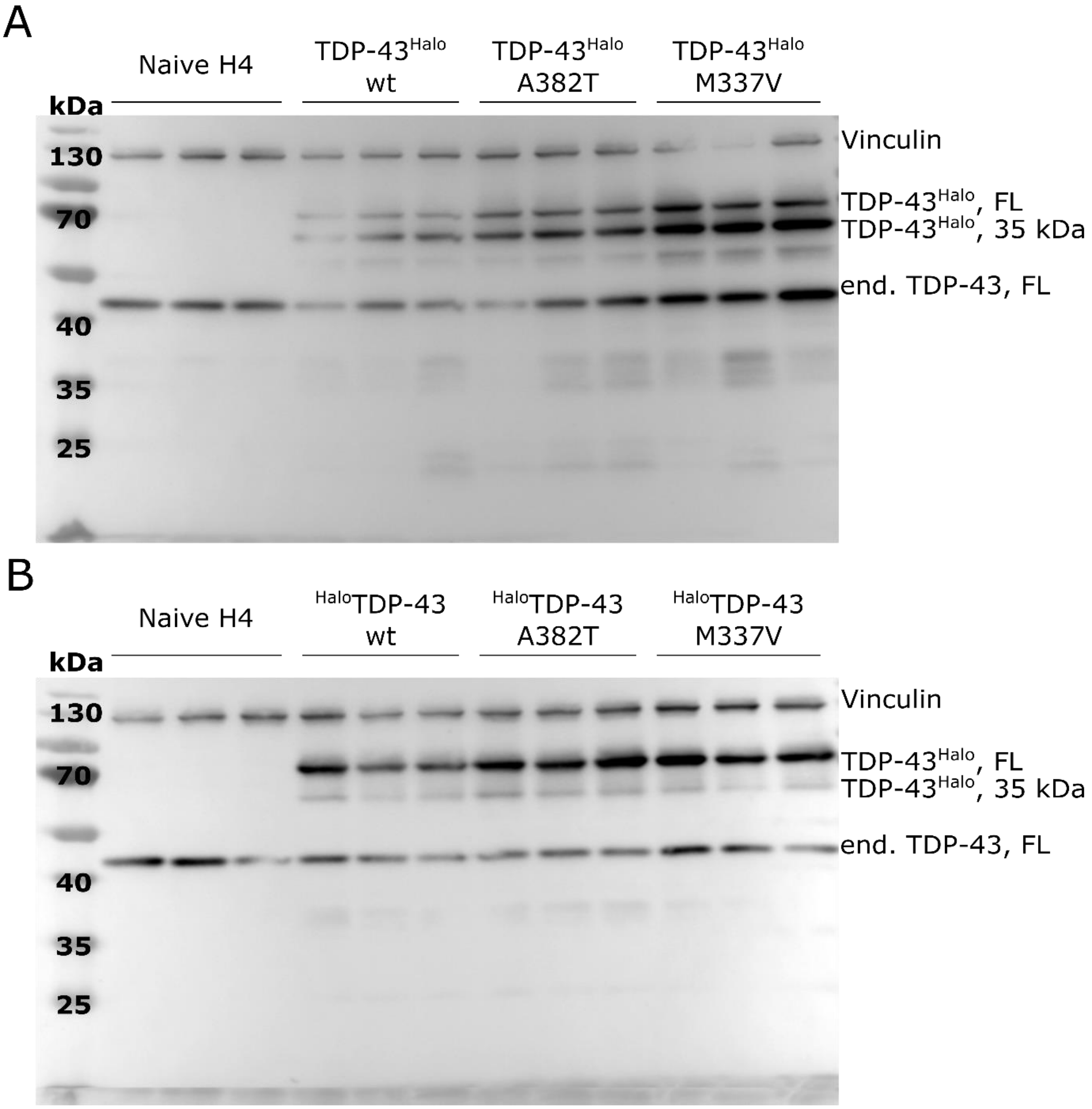


*Figure S2: Wester Blots for quantification of the overexpression of Halo-Tagged TDP-43 species. A. C-terminally tagged cell lines (wild-type, A382T, M337V) B. N-terminally-tagged cell lines (wild-type, A382T, M337V). Antibodies: anti-Vinculin, anti-TDP-43 (both 1:1000).*

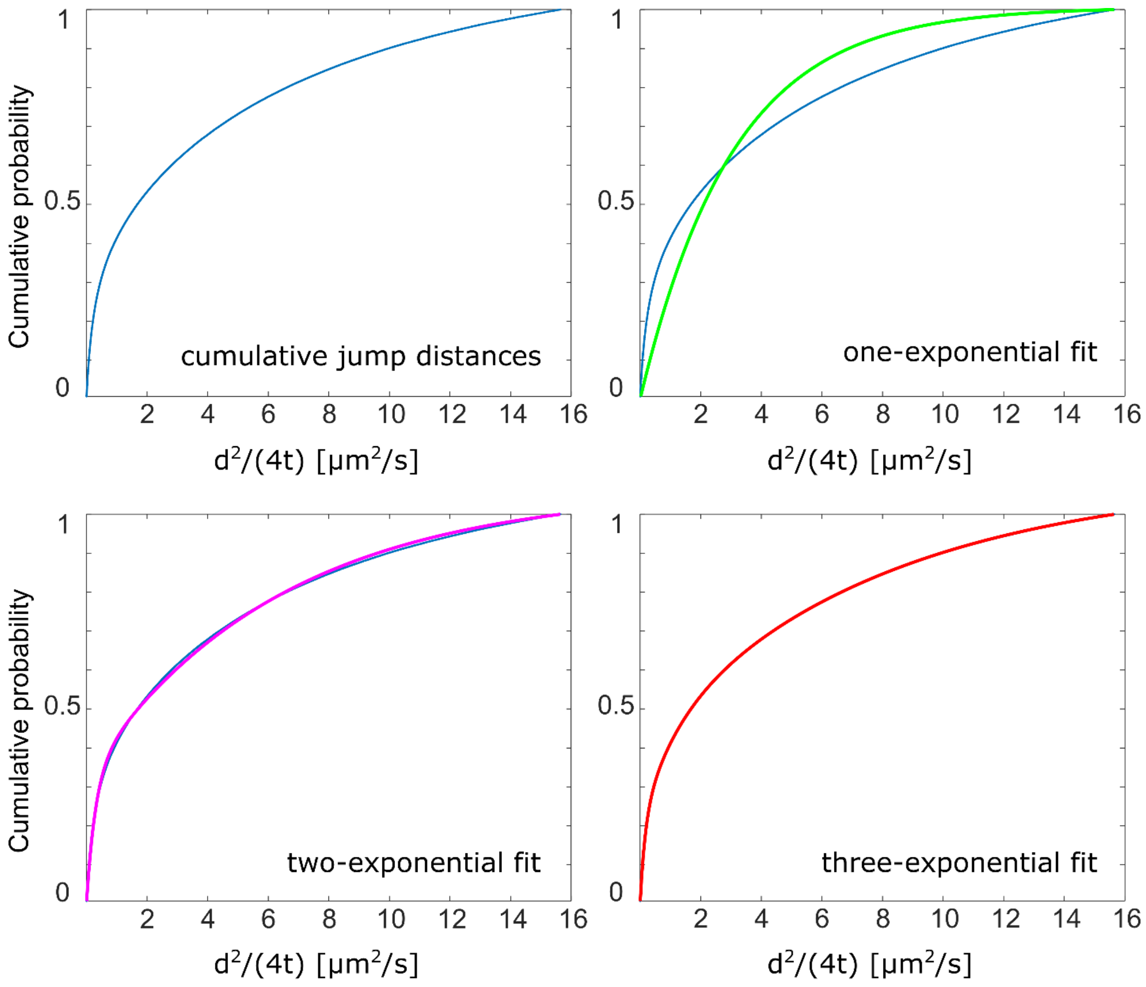


**Display Parameter**

| | |
|---|---|
| Bin number | 650 |
| Bin size [nm] | 1 |
| Pixel size [nm] | 0.13 |
| Jumps-to-consider | 5 |

**Fit Parameter - Start Values**

$D_1$ = 0.001
$D_2$ = 0.01
$D_3$ = 0.1

**SSE Values**

one-exponential fit = 4.615
two-exponential fit = 0.041
three-exponential fit = 0.001

**Adj. R squared values**

one-exponential fit = 0.914
two-exponential fit = 0.999
three-exponential fit = 1

*Figure S3: Assessment of fitting parameters for the multi-exponential fitting of cumulative distributions obtained from jump size histograms. Cumulative jump distance histogram (blue) and fitting curves overlays (1-rate exponential fit/green, 2-rate exponential fit/magenta, 3-rate exponential fit/red) and overview of fit errors, SSE (summed squared error) and adjusted $R^2$-error for all fitting conditions.*

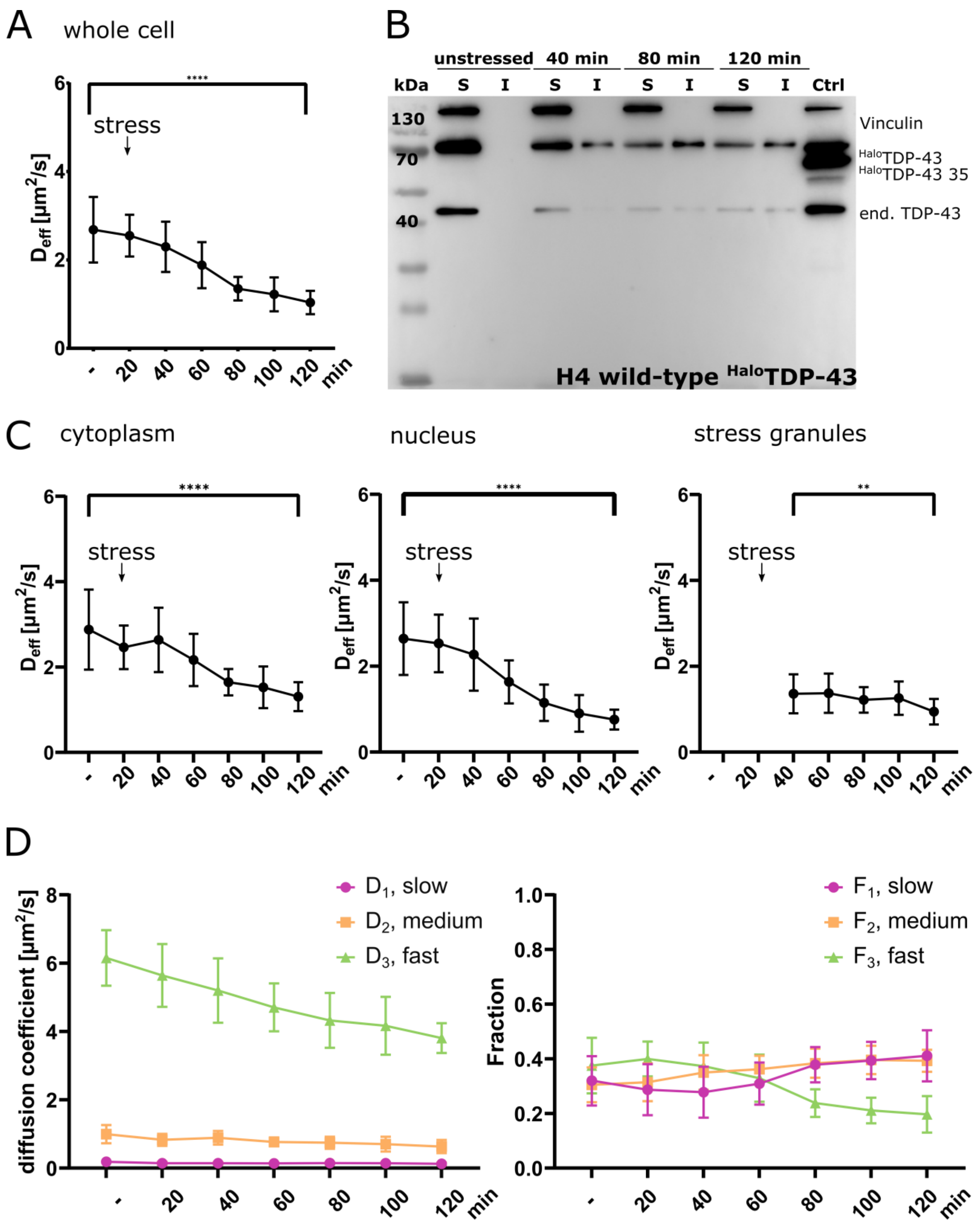


*Figure S4: Sodium arsenite stress leads to a reduction of TDP-43 mobility in specific cellular regions (cytoplasm, nucleus, stress granules). A. Stress time course of the effective diffusion*

*coefficient $D_{eff}$ for the $^{Halo}$TDP-43 construct (whole cell, mean + STD, Welch's t-test). B. Solubility Assay of the $^{Halo}$TDP-43 construct. Solubility was assessed under unstressed conditions and different stress time-points shows an increasing insoluble TDP-43 fraction with increasing stress duration (unstressed, 40 min, 80 min and 120 min of 0.5 mM sodium arsenite treatment, anti-Vinculin, anti-TDP-43). C. Stress time course of the effective diffusion coefficient $D_{eff}$ for the $^{Halo}$TDP-43 construct (cytoplasm, nucleus, stress granule, mean + STD, Welch's t-test). D. Analysis of the different diffusion constants (slow/$D_1$/magenta, medium/$D_2$/orange, fast/$D_3$/green) and the respective fraction within the stress time-course experiment (whole cell). For all experiments, standard deviations were calculated from the movie-wise distribution of the plotted value and statistical significance was assessed with a multiple unpaired t-test with Welch's correction. P-value ranges: <0.0001 ****, 0.0002 ***, 0.0021 **, 0.032 *, 0.123 ns.*

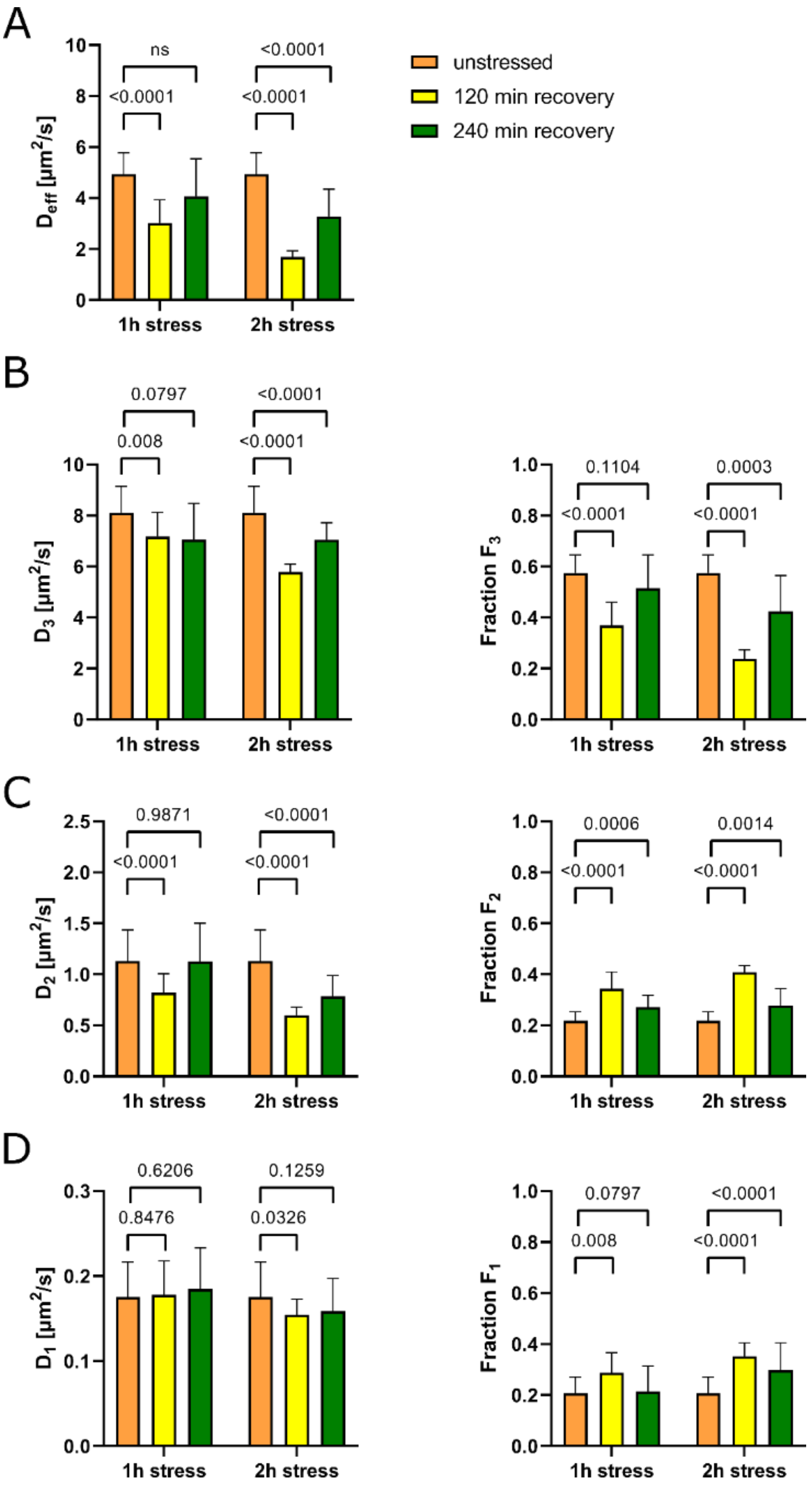
A
B
C
D
unstressed
120 min recovery
240 min recovery
$D_{eff}$ [µm²/s]
$D_3$ [µm²/s]
Fraction $F_3$
$D_2$ [µm²/s]
Fraction $F_2$
$D_1$ [µm²/s]
Fraction $F_1$
1h stress
2h stress
ns
<0.0001
0.0797
0.008
0.1104
0.0003
0.9871
0.0006
0.0014
0.6206
0.8476
0.1259
0.0326

*Figure S5: Statistical analysis of the effective diffusion constant $D_{eff}$ and the diffusion coefficients and fractions ($D_3/F_3$, $D_2/F_2$, $D_1/F_1$) of TDP-43 after 60 min or 120 min of sodium arsenite stress and 120 min or 240 min of recovery (orange: unstressed, yellow: 120 min recovery, green: 240 min recovery). A. Comparison of $D_{eff}$ between unstressed conditions and 120 min or 240 min of recovery after 60 min or 120 min of stress B – D. Comparison of $D_3/F_3$, $D_2/F_2$ and $D_1/F_1$ after 60 min or 120 min stress at unstressed condition, 120 min or 240 min of recovery. Statistical significance was assessed with a multiple unpaired t-test with Welch's correction. P-value ranges: <0.0001 ****, 0.0002 ***, 0.0021 **, 0.032 *, 0.123 ns.*

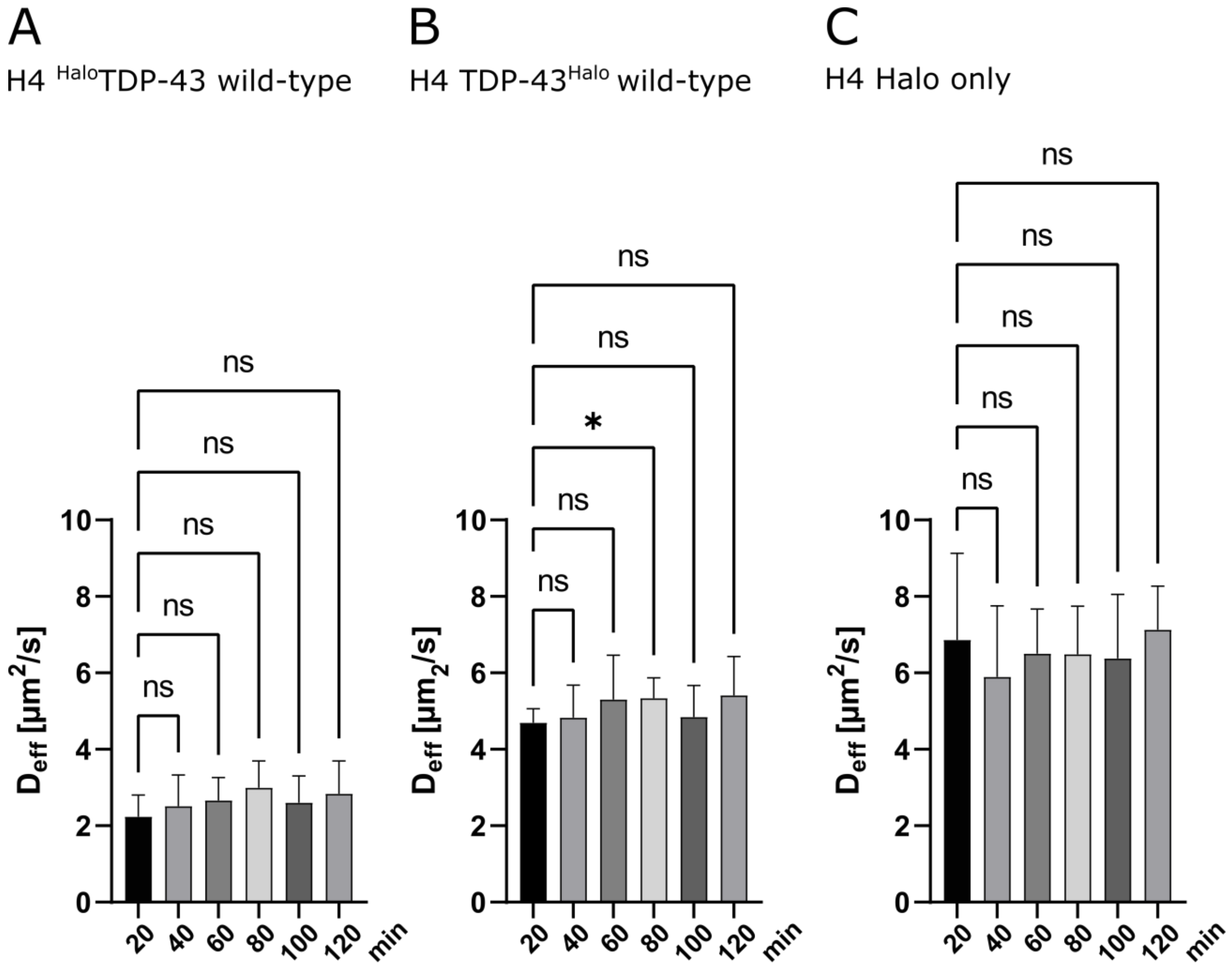


*Figure S6: Control measurements for single-molecule tracking measurements. A. Halo-only stress control. Mobility was assessed for 120 min under sodium arsenite stress. B and C. Setup control of C- and N-terminally tagged wild-type cell lines. Cells were imaged under unstressed conditions for 120 min to assess setup influences on TDP-43 mobility (Statistical test: Brown-Forsythe and Welch ANOVA).*

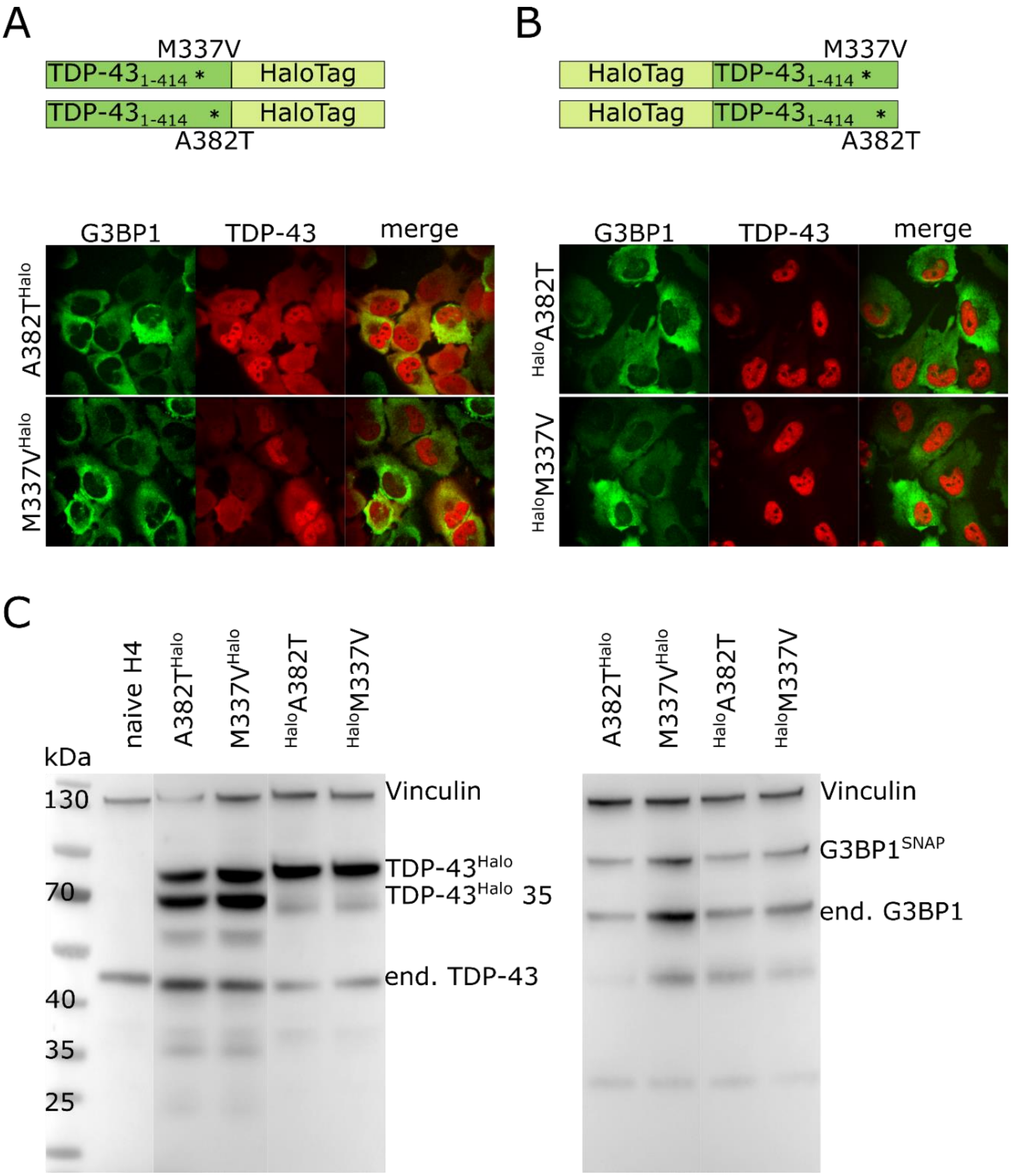


*Figure S7: Figure 1: Generation of mutant TDP-43 cell lines. A. Schematic overview of the used TDP-43$^{Halo}$ constructs and spinning disk confocal images of the double-transgenic (M337V$^{Halo}$, A382T$^{Halo}$) cell lines under unstressed conditions (red: TDP-43-TMR, green: G3BP-SiR, scale bar 10 µm). B. Schematic overview of the used $^{Halo}$TDP-43 constructs and spinning disk confocal images of the double-transgenic cell lines ($^{Halo}$M337V, $^{Halo}$A382T) under unstressed conditions*

*(red: TDP-43-TMR, green: G3BP-SiR, scale bar 10 µm). C. Western Blot overview of the double-transgenic cell lines stained with anti-vinculin, anti-TDP-43 or anti-G3BP1 antibodies.*

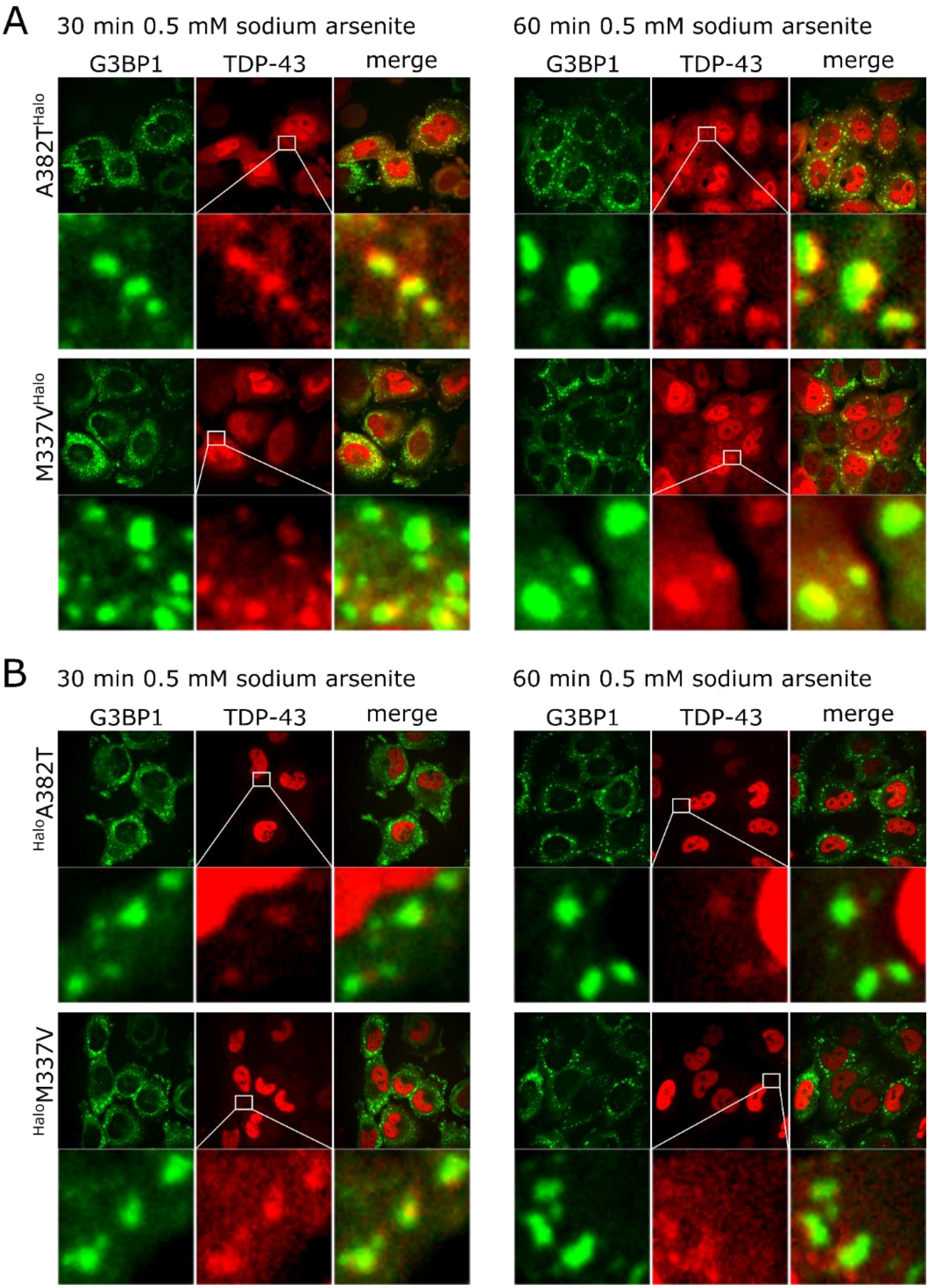
A
30 min 0.5 mM sodium arsenite
60 min 0.5 mM sodium arsenite
G3BP1
TDP-43
merge
A382T^Halo
M337V^Halo
B
30 min 0.5 mM sodium arsenite
60 min 0.5 mM sodium arsenite
G3BP1
TDP-43
merge
^HaloA382T
^HaloM337V

*Figure S8: Transgenic TDP-43 constructs are found in G3BP1-positive stress granules after 30 min and 60 min of 0.5 mM sodium arsenite treatment. A. Spinning disk confocal images of all for all C-terminally tagged TDP-43 constructs (A382T$^{Halo}$ and M337V$^{Halo}$ mutants) after 30 min and 60 min 0.5 mM sodium arsenite treatment. Exemplary stress granules are marked with an rectangle. B. Spinning disk confocal images of all for all N-terminally tagged TDP-43 constructs ($^{Halo}$A382T and $^{Halo}$M337V mutants) after 30 min and 60 min 0.5 mM sodium arsenite treatment. TDP-43 granules are depicted with white rectangle (red: TDP-43-TMR, green: G3BP-SiR, scale bar 10 µm).*

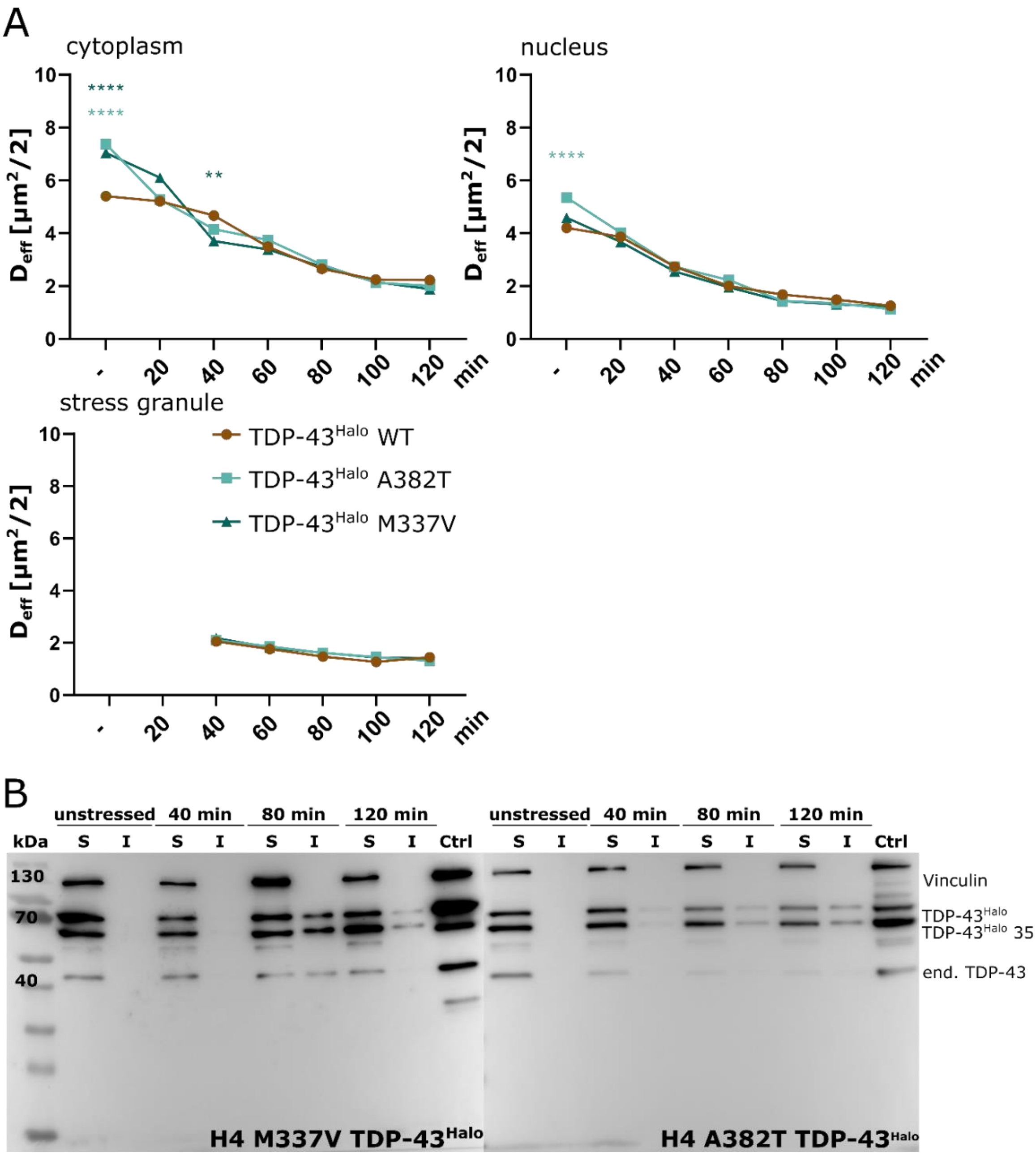


*Figure S9: Region-wise TDP-43 mobility assessment and comparison between mutant and wild-type cell lines. A. Comparison of the stress related TDP-43 mobility of C-terminally taggedTDP-43 wild-type and mutant (M337V, A382T) constructs in the cytoplasm, nucleus and stress granules. Comparison shows statistical significance between the wild-type and the mutant TDP-43 at early stress conditions but no further differences at later time-points. B. Solubility assay of the C-terminally tagged TDP-43 mutants (M337V, A382T) for unstressed and stress conditions (40 min, 80 min, 120 min). Solubility assessment showed an increased insoluble fraction of both mutant TDP-43 constructs with increasing stress duration (anti-vinculin, anti-TDP-43).*

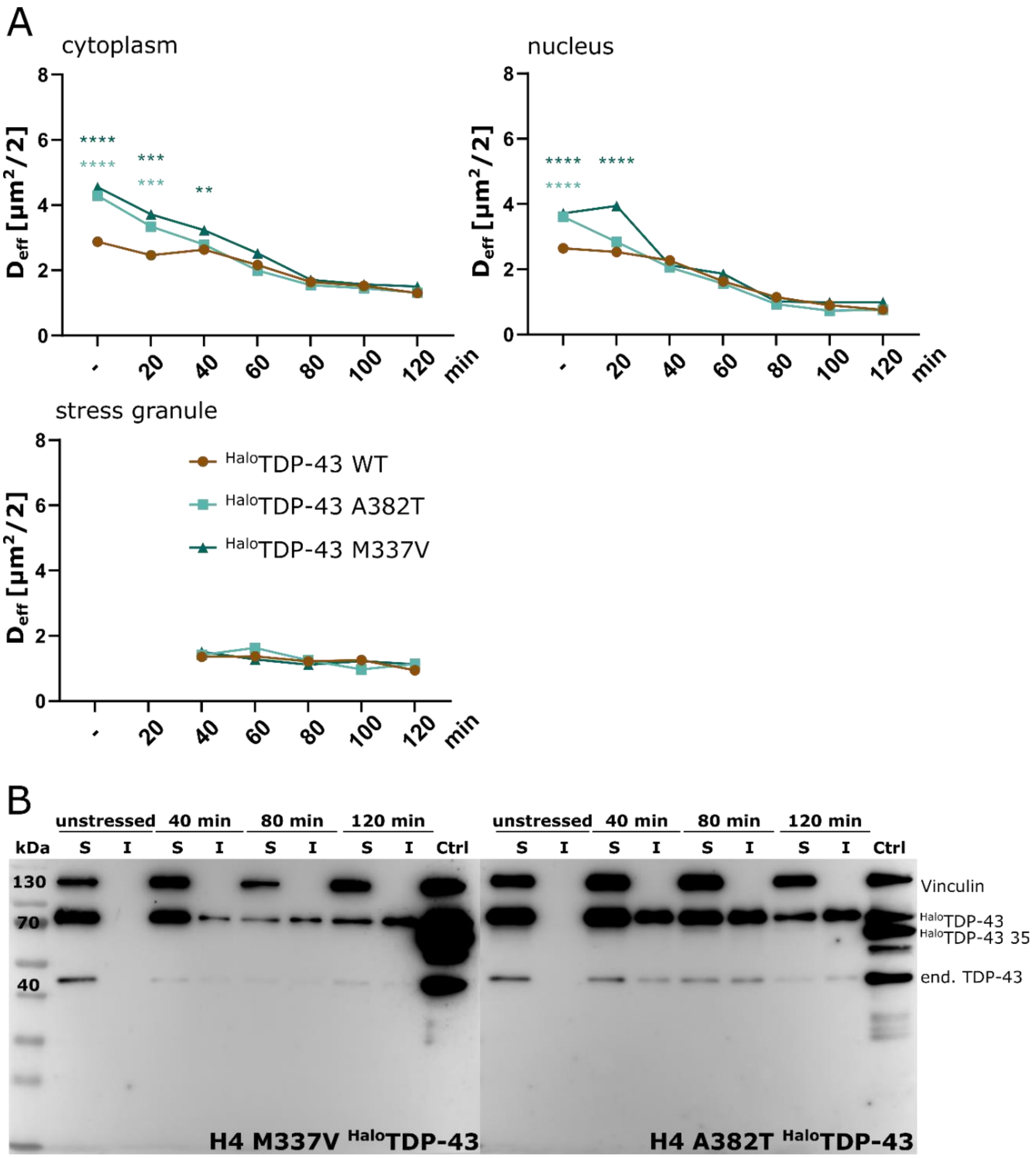


*Figure S8: Region-wise TDP-43 mobility assessment and comparison between mutant and wild-type cell lines. A. Comparison of the stress related TDP-43 mobility of N-terminally taggedTDP-43 wild-type and mutant (M337V, A382T) constructs in the cytoplasm, nucleus and stress granules. Comparison shows statistical significance between the wild-type and the mutant TDP-43 at early stress conditions but no further differences at later time-points. B. Solubility assay of the C-terminally tagged TDP-43 mutants (M337V, A382T) for unstressed and stress conditions (40 min, 80 min, 120 min). Solubility assessment showed an increased insoluble fraction of both mutant TDP-43 constructs with increasing stress duration (anti-vinculin, anti-TDP-43)*

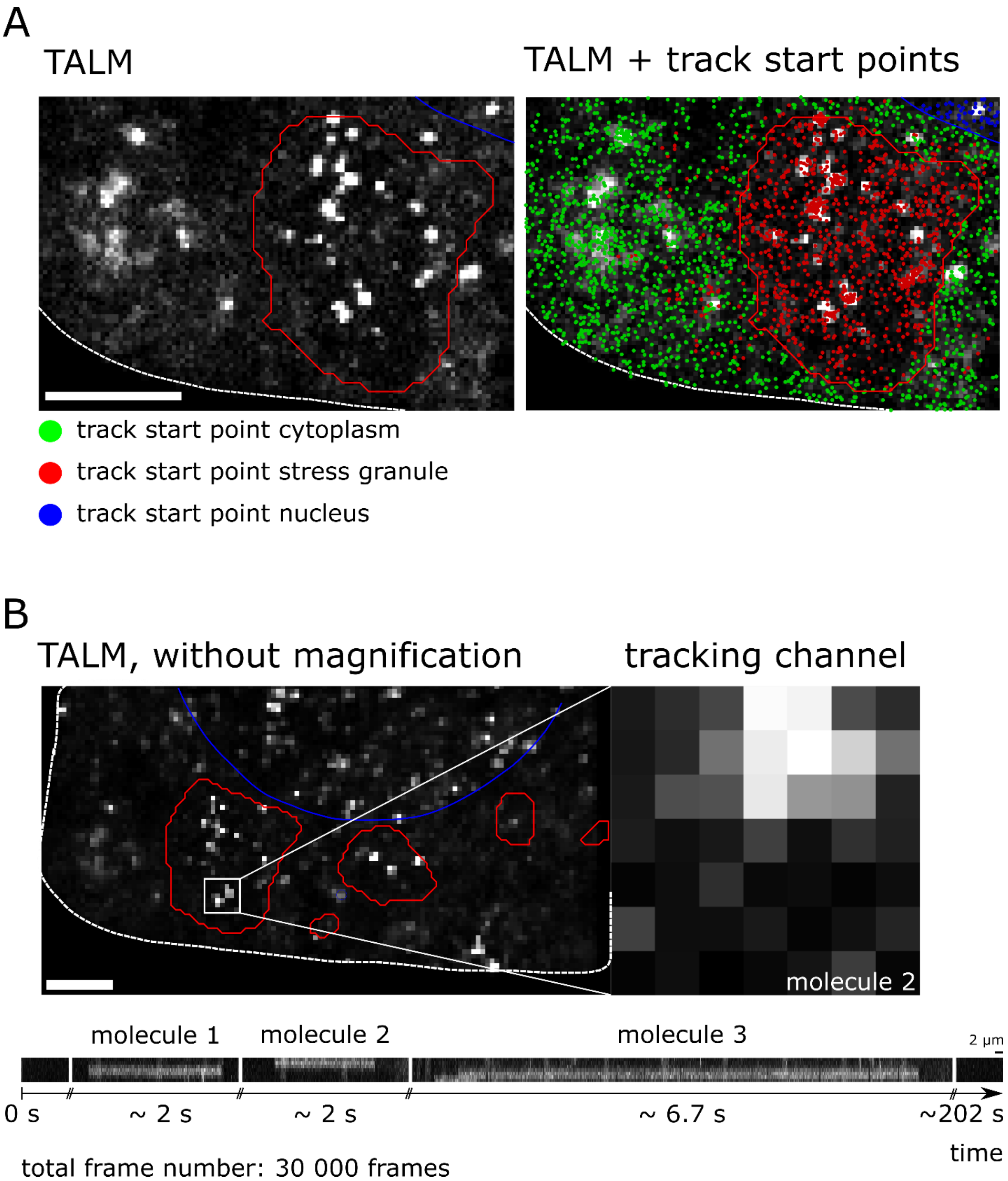


*Figure S9: A. Track start point overlay with TALM images (cytoplasm: green, stress granule: red, nucleus: blue, scale bar 2 µm). B. Kymograph analysis of binding hotspots within stress granules shows repeated binding of TDP-43$^{Halo}$ to the same region (scale bar 2 µm). Note that the displayed time-spans show localizations which are separated by much longer time periods without localizations, indicating single-molecule conditions.*
